# Morpho-functional characterization of the endo-lysosomal system by high-throughput correlative light-electron microscopy

**DOI:** 10.1101/2021.05.21.445146

**Authors:** Jan van der Beek, Cecilia de Heus, Nalan Liv, Judith Klumperman

## Abstract

Rab5, EEA1 and APPL1 are frequently used in fluorescence microscopy to mark early endosomes, whereas Rab7 is used as marker for late endosomes and lysosomes. However, since these proteins localize poorly in immuno-electron microscopy, systematic studies on their ultrastructural distributions are lacking. Here we address this gap by presenting a quantitative, high-throughput, on-section correlative light-electron microscopy (CLEM) approach using the sensitivity of fluorescence microscopy to infer label to hundreds of organelles classified by ultrastructure. We show that Rab5 predominantly marks small, endocytic vesicles and early endosomes. EEA1 co-localizes with Rab5 on especially early endosomes, but unexpectedly also labels Rab5-negative late endosomes and even lysosomes. APPL1 is restricted to small Rab5-positive, vesicular profiles without any visible content or ultrastructural marks. Rab7 primarily labels late endosomes and lysosomes. Our studies reveal the first ultrastructural distribution of key endosomal proteins at their endogenous levels and introduce CLEM as sensitive alternative for quantitative immuno-EM.

## Introduction

A ubiquitous feature of eukaryotic cells is the division of labor over distinct functional compartments. The endo-lysosomal system contains different compartments, which together define the ultimate fate of internalized and internal molecules. Mutations in endo-lysosomal proteins cause severe storage disorders^1^ and disorganization of the endo-lysosomal system is an underlying cause in cancer, neurological conditions and many other diseases^2–6^. Understanding changes in the endo-lysosomal system in relation to cellular physiology is therefore a topic of intense research and a fundamental step in elucidating human pathologies.

Endo-lysosomal compartments are functionally distinguished by their capacity for cargo sorting, recycling and degradation and, more recently, transcriptional signaling to the nucleus^7^. Following endocytosis from the plasma membrane, early endosomes uncouple ligands from receptors and sort proteins for recycling or degradation^8–10^. Early endosomes mature into late endosomes^11–13^, which recycle proteins to the Trans Golgi Network (TGN)^10,14^ and are capable of fusion with autophagosomes and lysosomes^12,15,16^. Lysosomes are the compartments with the highest level of active hydrolases that break down the enclosed content, providing nutrients and new building blocks for the cell. Late endosomes and lysosomes sense the cells’ nutrient status and signal this to the nucleus to regulate the transcription of lysosome-and autophagy-related genes^7,17,18^. Together, this highly interconnected and dynamic system of organelles determines protein turnover and maintains cellular homeostasis.

The different endo-lysosomal compartments are defined by stage-specific molecular machinery and morphological characteristics^19,20^. Small GTPases are the master regulators of membrane trafficking and, together with their effector proteins, mediate fusion, fission, trafficking, and signaling^7,10,21–31^. The small GTPase Rab5 is recruited to newly formed endocytic vesicles and early endosomes^25,32,33^, marking the early stages of endocytosis committed to recycling and sorting. Rab5-positive membranes form two subpopulations by attracting different effector proteins: ‘Adaptor protein, phosphotyrosine interacting with PH domain and leucine zipper 1’ (APPL1) and ‘Early Endosome Antigen 1’ (EEA1)^34–36^. APPL1 is a multifunctional adaptor protein forming a scaffold for a variety of signaling proteins^37^ and marks endosomes with a high propensity for fast recycling^34^. The long coiled-coil tether EEA1 enacts fusion between Rab5-positive vesicles and early endosomal vacuoles^28^. EEA1 remains present on maturing early endosomes^38^ till a change from Rab5 to Rab7 occurs^8^ that is driven by the Ccz1-Mon1 complex^12,13^. Rab7 activates numerous effector proteins, including Retromer for retrograde trafficking and the HOPS tethering complex^14,26,39,40^ required for late endosome – lysosome fusion.

The morphology of endosomes and lysosomes has been studied for decades using different types of Electron Microscopy (EM) methods. This has revealed essential structure-function relationships at the nanometer scale^20^. In general, tubules and clathrin coats are associated with sorting and recycling of cargo’s^9,41–45^, while intraluminal vesicles (ILV’s) and dense content are linked to the degradative pathway^46–49^. In addition, EM has revealed essential information on cellular context, such as type and number of contact sites of endo-lysosomes with ER and mitochondria^50–54^. Furthermore, immuno-EM methods have been instrumental in localizing proteins to the distinct endosomal sub-domains, such as recycling tubules, clathrin coats or ILV’s^41,42,44,55–60^. Collectively, these EM studies have provided an integrated view on the function, molecular composition and morphology of the different endo-lysosomal compartments and their subdomains^20^.

Because of their central roles in the endo-lysosomal system, Rab5, Rab7, EEA1 and APPL1 are topic of numerous studies. Moreover, Rab5 and EEA1 are frequently used in fluorescence microscopy to mark early endosomes, whereas Rab7 is a commonly-used marker for late endosomes and lysosomes^13,34^. However, the ultrastructural localization of these proteins has proven difficult and only few studies are reported. Using immuno-EM on thawed cryosections, EEA1^61,62^ has been localized to early endosomal vacuoles and overexpressed Rab7-GFP to late endosomes, lysosomes and autophagosomes^63,64^. APPL1 has been detected on tubular endosomes using pre-embedding labeling and silver-enhancement^34^, as well as through immuno-EM using a non-commercial antibody^35^. Using super-resolution correlative light-electron microscopy (CLEM) on 250 nm cryosections, Frank and colleagues localized Rab5-GFP to restricted domains of early endosomal vacuoles^65^. However, none of these approaches included a systematic, quantitative analysis of the ultrastructural distribution of these proteins. Nor has simultaneous labeling of multiple markers using the same system and methodology been performed. Moreover, the use of over-expression approaches may induce artefacts in endo-lysosomal morphology and lead to non-specific membrane associations^63^. Thus, a robust, quantitative ultrastructural analysis of organelles that are Rab5, Rab7, EEA1 or APPL1-positive is currently lacking. Additionally, it remains unknown how their distribution relates to the commonly used morphological definitions of endo-lysosomal organelles used in EM studies.

To connect functional-molecular information to morphology, we here present a high-throughput CLEM approach based on the use of ultrathin cryosections. Using optimized strategies for correlation, we detect the endosomal marker proteins by fluorescence microscopy, and then image the same sample in EM for accurate correlation of fluorescence labeling to ultrastructure^66,67^. We enable the correlation of hundreds of fluorescent spots to endo-lysosomal morphology, followed by a systematic categorization based on ultrastructure. Our data show that Rab5 mostly marks endocytic vesicles and early endosomes, and only partially overlaps with EEA1, which marks early endosomes and shows a surprisingly high level of association with late endosomes. Moreover, we observe that APPL1/Rab5 positive organelles are small, vesicular endosomes, which are clearly different from EEA1 compartments. Rab7 marks late endosomes and lysosomes displaying morphological elements indicative of degradation. Our studies introduce CLEM as a quantitative protein localization method that is a feasible and attractive high-throughput addition to ‘classical’ immuno-EM methods and provides novel ultrastructural insights on the distribution of commonly used endo-lysosomal markers.

## Results

### IF of endogenous Rab5, Rab7, APPL1 and EEA1 reveals distinct organelle populations

We selected a panel of commercially available antibodies against Rab5, Rab7, APPL1 and EEA1 that have been successfully used for Immuno-Fluorescence (IF) labeling studies (see **Fig**. 1, **Table** 2 in materials and methods). We first tested these antibodies in a conventional IF protocol on permeabilized HeLa cells fixed in 4% formaldehyde (FA). By double-labeling we addressed the overlap between the different proteins (**Fig**. 1). We found that Rab5-positive endosomes are present in both the cell periphery and in the perinuclear area (**Fig**. 1A, D, E), whereas Rab7-positive compartments are enriched in the perinuclear area (**Fig**. 1A-C). Due to the efficiency of Rab5 to Rab7 conversion by the Mon1-Ccz1 complex^12^ few endosomes with both Rab5 and Rab7 are expected^12,13^. Indeed, by IF (**Fig**. 1A) circa 6% of the Rab5-positive spots also labeled for Rab7. Vice versa, 24% of the Rab7-positive spots was also positive for Rab5 (**Fig**. 1G). The rest of Rab5 or Rab7 spots formed separate pools. Rab5-positive spots largely overlapped with EEA1 and APPL1 staining (**Fig**. 1D, E, I), whereas Rab7-positive spots showed 35% and 22% overlap with the late endosomal/lysosomal markers^68,69^ CD63 and Cathepsin D, respectively (**Fig**. 1B, C, H). The finding that a sizeable portion of Rab7-positive compartments does not contain CD63 or Cathepsin D is somewhat unexpected but has several putative explanations. First, the spatial segregation between luminal CD63 and Cathepsin D and membrane-associated Rab7 may decrease the level of co-localization (**Fig**. 1H). Second, Rab7 in addition to endosomes can also be found on small vesicles, reported before^14^ and seen later in **figures** 3D, 5C and 5D. And finally, CD63 and Cathepsin D may only label a subset of endosomes and lysosomes.

**Figure 1.**
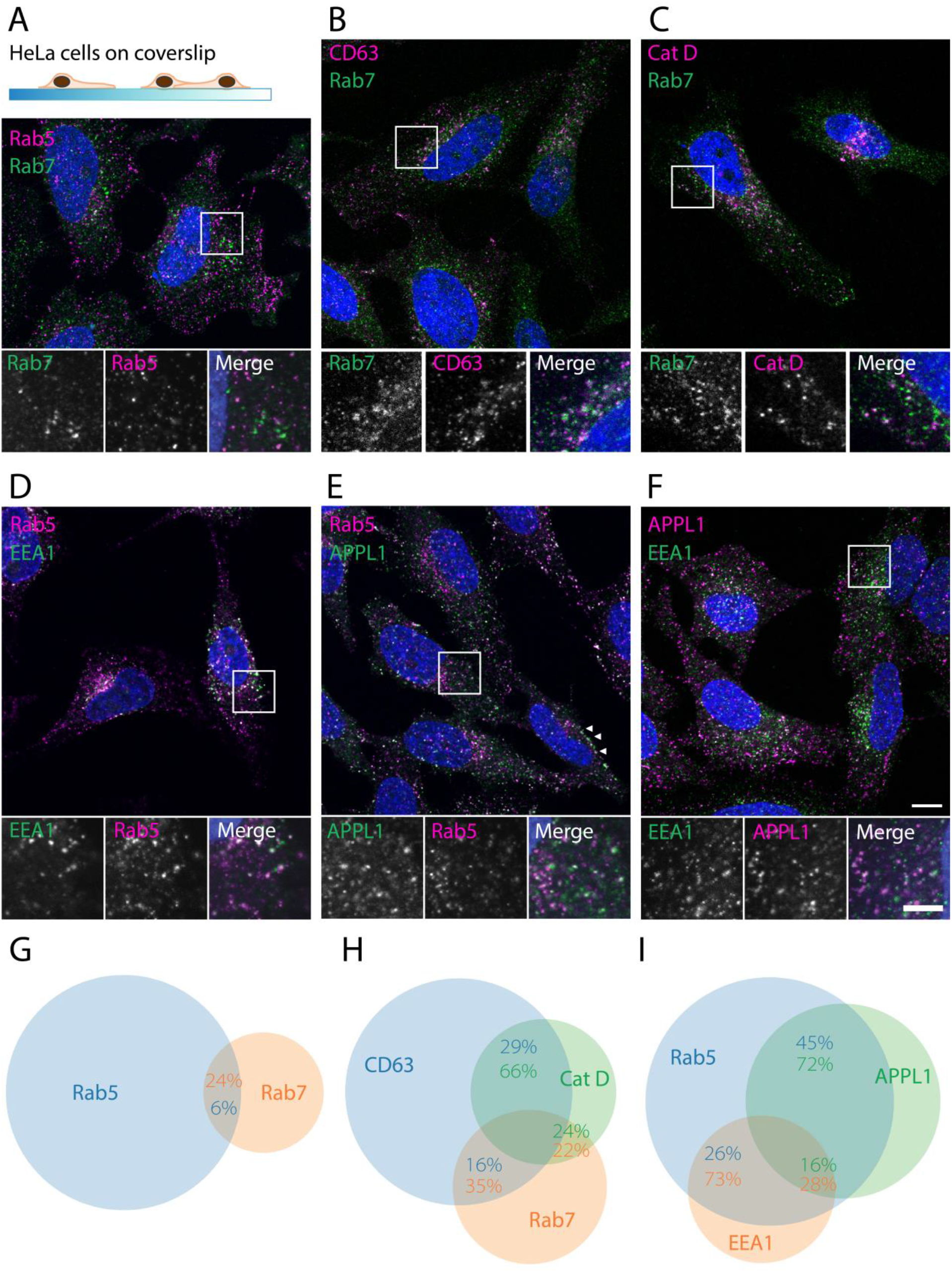
Immuno-fluorescence of endo-lysosomal markers reveals overlapping yet separate localization patterns. Pictures are confocal images (slices) of double-labeled, permeabilized HeLa cells fixed with 4% FA. A: Rab5 and Rab7 predominantly mark separate organelles. B, C: Rab7 co-localizes partially with CD63 and Cathepsin D. D: EEA1 labels a perinuclear subpopulation of Rab5 endosomes. E: APPL1 labels a peripheral pool of Rab5 endosomes. Note the presence of APPL1 endosomes just below the plasma membrane (arrowheads). F: EEA1 and APPL1 show very little overlap. G-I: Venn diagrams based on co-localization analysis of labeling combinations in A-F. Circle size is proportional to total dots detected for a protein, overlap to number of co-localized dots. Images were analyzed by dot detection in two or three channels, after which overlapping dots were classified as co-localized particles. Percentages represent the co-localized fraction of the correspondingly colored protein. See methods for a more detailed description of the analysis, see Table S1 for standard deviations, cell and organelle numbers. Scale bars 10µm in larger images, 5µm in insets.

**Table 1.** Effect of fixation on immuno-fluorescence intensity and EM morphology. 90 nm Cryosections of HeLa cells fixed according the indicated protocols were fluorescently labeled and imaged using fixed settings. The signal-to-noise-ratio (SNR) was calculated as mean intensity value of the 0.5% brightest pixels divided by the mean intensity value of the reverse selection. 5 Field-of-views were averaged for each measurement. Most SNRs significantly decline upon >30 minutes FA fixation or addition of GA. Morphological quality of EM images was based on blind ranking by 6 experienced electron microscopists. Based on these measurements 1 hour fixation with 4% FA was chosen as best fixative for CLEM. FA = Formaldehyde, GA = Glutaraldehyde, ON = Overnight.

| **Fixation protocol** | **IF SNR Rab5** | **IF SNR Rab7** | **IF SNR EEA1** | **IF SNR APPL1** | **EM morphology** | **Supp. Figure** |
| --- | --- | --- | --- | --- | --- | --- |
| 30min FA 4% | 2,09 ±0,05 | 1,61 ±0,03 | 2,00 ±0,06 | 2,30 ±0,11 | - | S1A |
| 1h FA 4% | 1,72 ±0,15 | 1,66 ±0,07 | 1,76 ±0,08 | 1,85 ±0,06 | +/- | S1B |
| 2h FA 4% | 1,62 ±0,34 | 1,51 ±0,03 | 1,42 ±0,04 | 1,63 ±0,03 | +/- | S1C |
| 1h FA 4% → |  |  |  |  | + | S1D |
| ON FA 0.6% | 1,44 ±0,07 | 1,48 ±0,06 | 1,41 ±0,04 | 1,66 ±0,06 |  |  |
| ON FA 4% | 1,43 ±0,13 | 1,51 ±0,02 | 1,24 ±0,02 | 1,31 ±0,02 | + | S2A |
| 30min FA 4% |  |  |  |  | ++ | S2B |
| + GA 0.2% | 1,66 ±0,06 | 1,36 ±0,03 | 1,20 ±0,04 | 1,28 ±0,08 |  |  |
| 2h FA 4% + |  |  |  |  | ++ | S2C |
| GA 0.2% | 1,70 ±0,02 | 1,41 ±0,03 | 1,34 ±0,05 | 1,39 ±0,04 |  |  |
| ON FA 4% + |  |  |  |  | ++ | S2D |
| GA 0.2% | 1,64 ±0,04 | 1,36 ±0,02 | 1,25 ±0,01 | 1,30 ±0,02 |  |  |

**Table 2.**
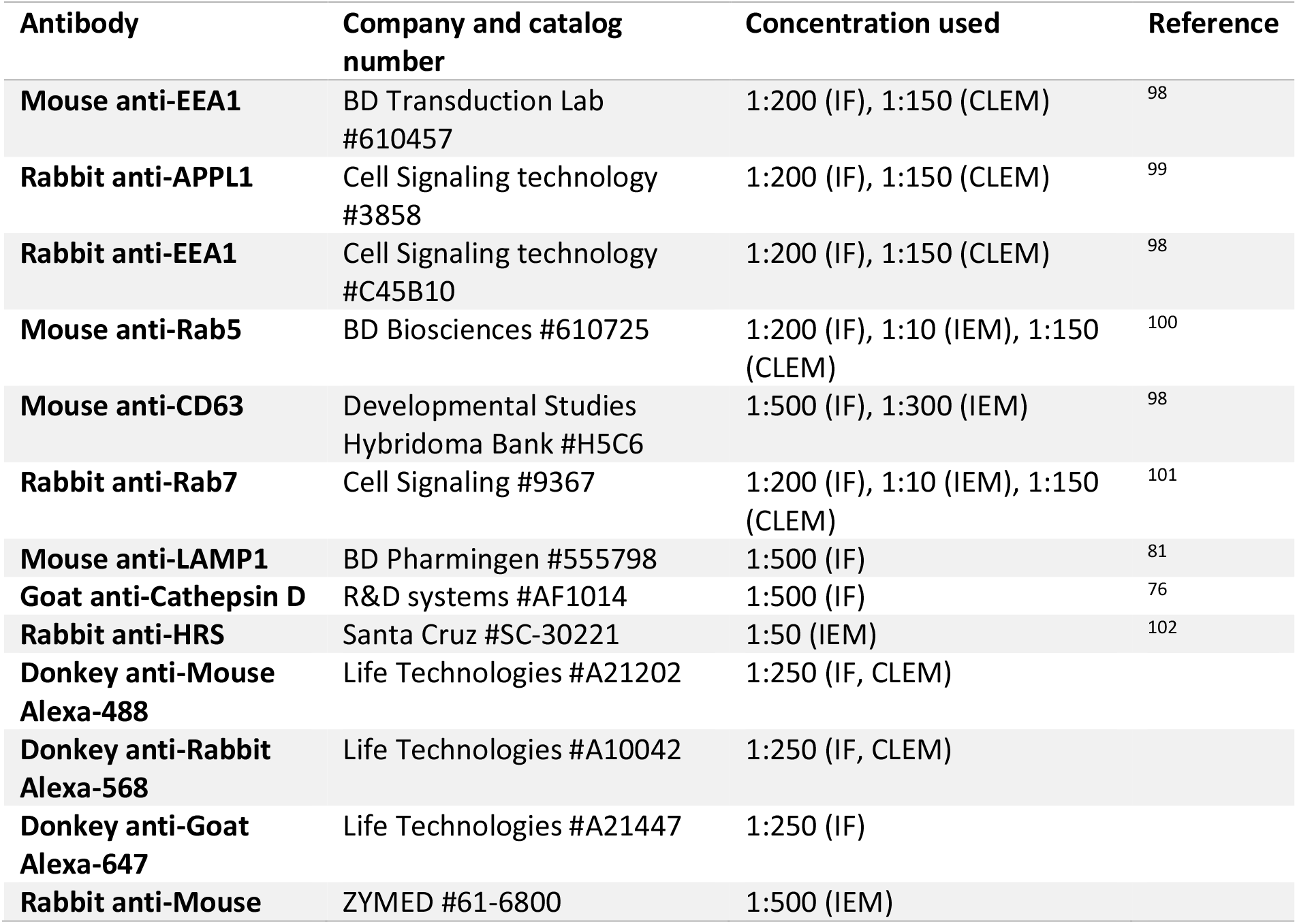
Antibodies used in this study. IF, Immuno-Fluorescence; CLEM, Correlative Light-Electron Microscopy; IEM, Immuno-EM.

**Figure 2.**
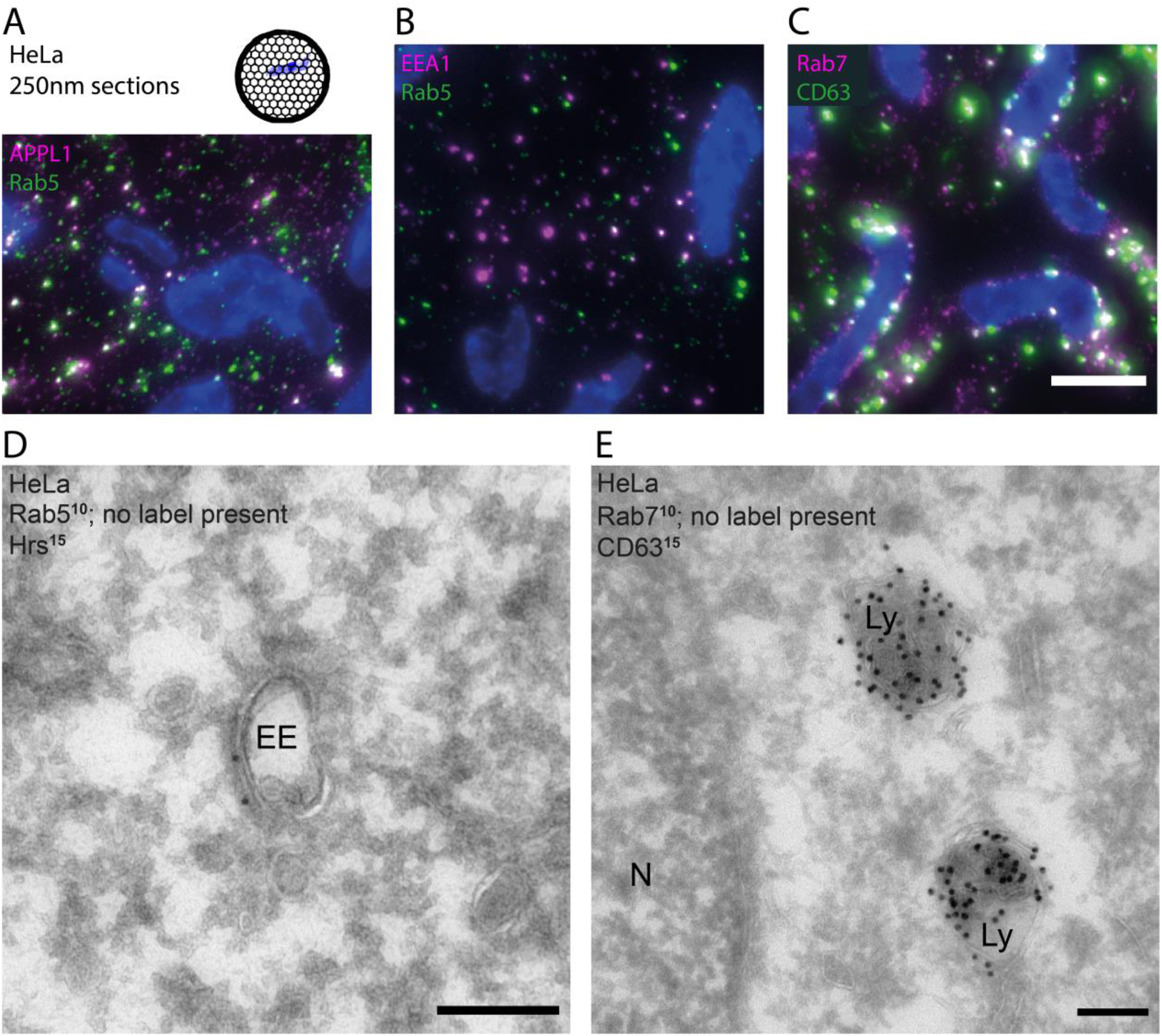
Immuno-staining of Rab5 and Rab7 on cryosections yields fluorescent but no immunogold labeling. HeLa cells fixed with 4% FA for 1 hour and processed into (A-C) 250 nm cryosections and imaged by widefield microscopy or (D-E) 90 nm cryosections and imaged by TEM. A, B, C: On-section immuno-labeling for Rab5, Rab7, CD63, EEA1 and APPL1 in indicated combinations using the same primary antibodies as in Figure 1. Scale bar 5 µm. D, E: Double-immunogold labeling using our established immuno-EM protocol^72^ with the same primary antibodies as in A-C, subsequently labeled with protein-A conjugated to gold particles. Gold particle size indicated in superscript. D: Example of early endosome (EE) with as positive control immuno-EM localization of the early endosome-associated protein Hrs (15 nm gold). However, Rab5 label (10 nm gold) could not be detected by immuno-EM. E: Example of two lysosomes (Ly) abundantly labeled for the late endosomal/lysosomal protein CD63 (15 nm gold) labeling. Rab7 label (10 nm gold) could not be found by this immuno-EM approach. Scale bars 200 nm.

**Figure 3.**
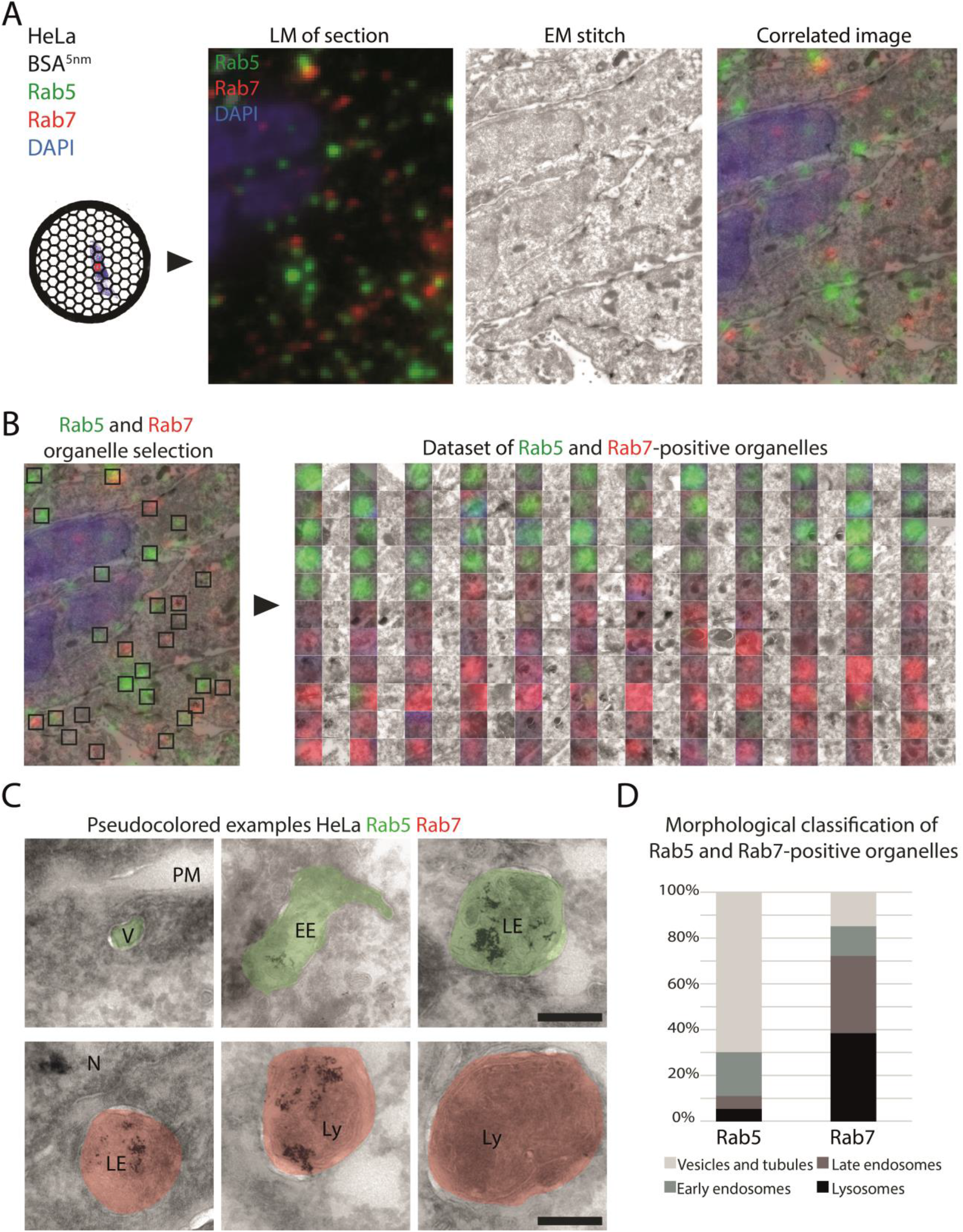
High-throughput CLEM of Rab5 and Rab7 reveals complementary distributions over early and late endo-lysosomal compartments. CLEM of Hela cells fixed with 4% FA for 1 hour. Prior to fixation cells were incubated with BSA-gold^5nm^ for 3h. A: Left: Widefield image of part of a 90 nm cryosection labeled for Rab5 and Rab7 and AlexaFluor488 and -568 secondary antibodies, respectively. Middle: Stitched EM image of the same area composed of 63 43,000x magnification images. Right: overlay of IF and EM images. B: Left: Low magnification overview of organelles selected for correlation (indicated by black squares). Right: High-throughput dataset of IF to EM-correlated Rab5- and Rab7-positive organelles. Each left row shows the overlay image with at the right the EM only. C: Zoom-ins of pseudocolored examples of Rab5 (green) and Rab7 (red) positive organelles similar as shown in B. Note that some organelles contain internalized BSA-gold^5nm^. For original images see Fig. S4A. D: Relative distribution of Rab5 and Rab7 over distinct endo-lysosomal compartments. N = 37 for Rab5 and 64 for Rab7, taken from 3 double-labeled samples. EE; early endosome, LE; late endosome, Ly; lysosome, N; nucleus, PM; plasma membrane, V; vesicle. Scale bars 200 nm.

**Figure 4.**
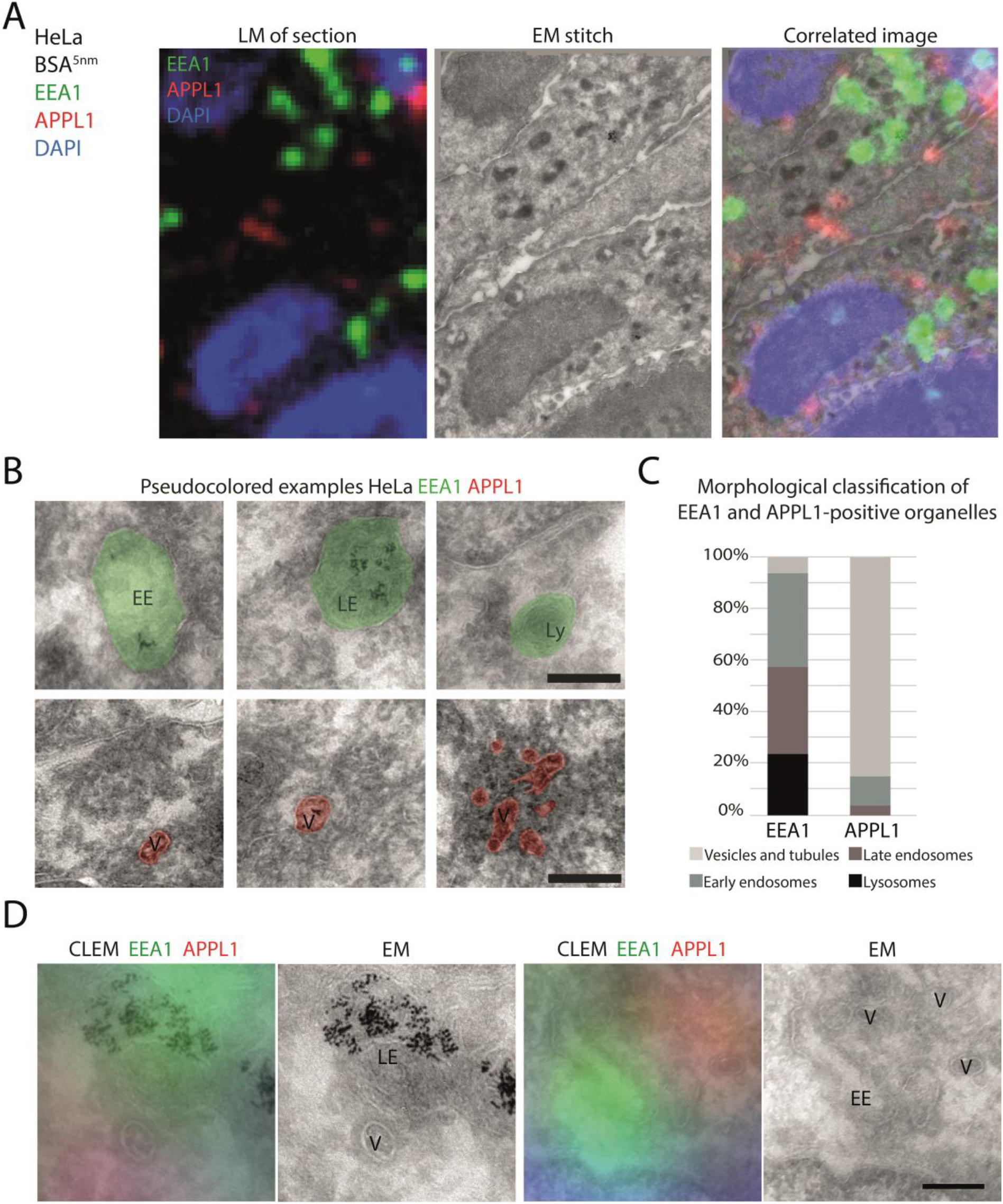
High-throughput CLEM of EEA1 and APPL1 shows strikingly distinct distributions. A-C: HeLa cells prepared as in Figure 3; labeled for EEA1 and APPL1 with AlexaFluor488 and -568, respectively. For original images of B, see Fig. S4B. C: Relative distribution of EEA1 and APPL1 over distinct endo-lysosomal compartments. N = 84 for EEA1 and 54 for APPL1, taken from 3 different samples. EEA1 predominantly labels early and late endosomes. APPL1 is almost exclusively found on small vesicles. D: Co-localization of EEA1 and APPL1 is rare. Fluorescent spots showing both labels by CLEM appear as early endosomes with a nearby vesicle. EE; early endosome, LE; late endosome, Ly; lysosome, V; vesicle. Scale bars 200 nm.

**Figure 5.**
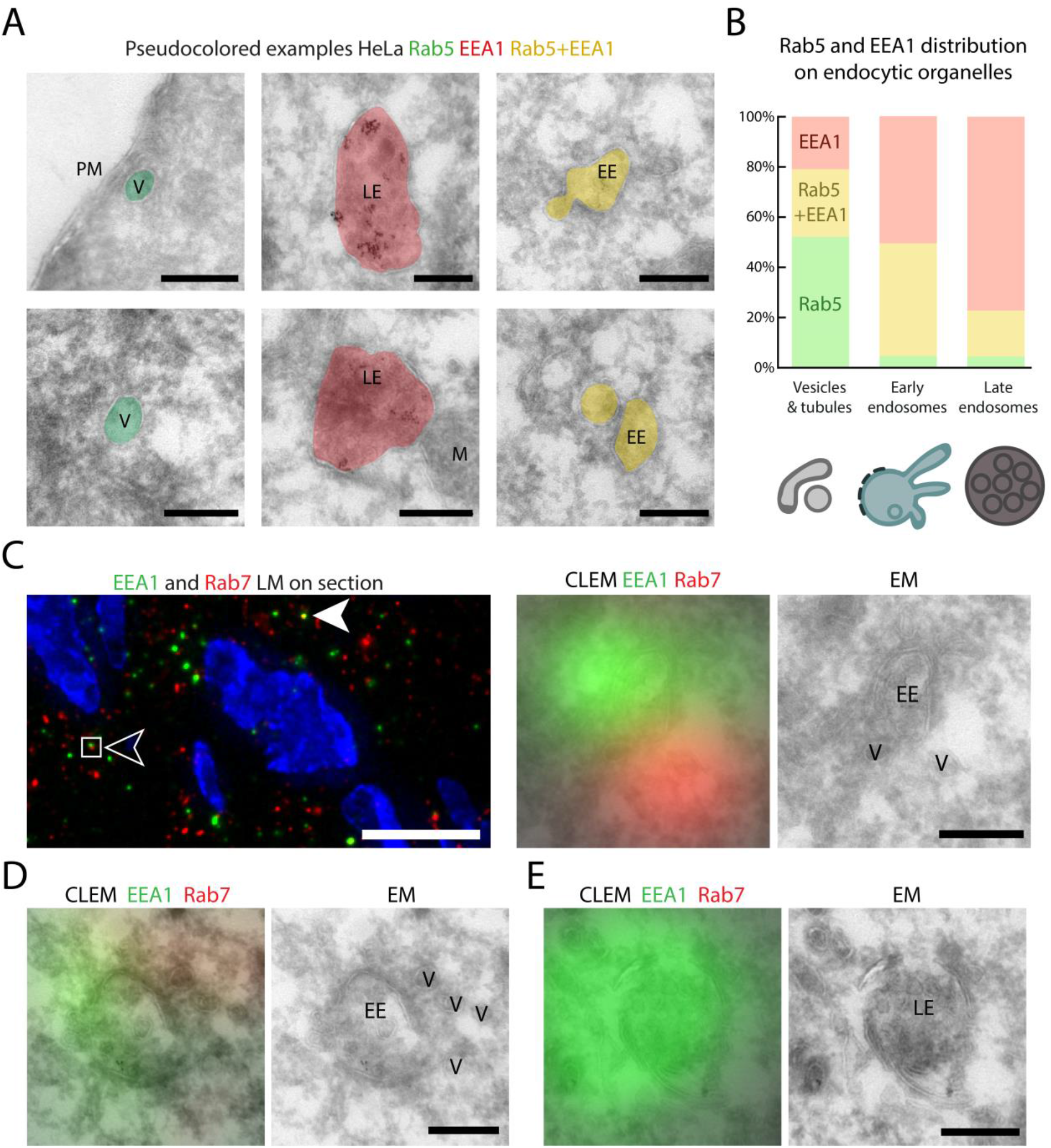
EEA1 localizes to late endosomal compartments that lack Rab5. HeLa cells prepared for CLEM as in Figure 3. A: Pseudocolored CLEM of Rab5 and EEA1 labeled with AlexaFluor488 and -568. Exclusive Rab5 or EEA1, or double-labeling is represented by green, red or yellow pseudocoloring, respectively. Original EM images are shown in Fig. S5. B: Morphological classification of Rab5, Rab5 + EEA1 and EEA1-positive organelles. N = 40, 62 and 76 for Rab5, Rab5 + EEA1 and EEA1, respectively. Data collected from 3 different samples. C, left: IF of HeLa cryosections labeled for EEA1 and Rab7 with AlexaFluor488 and - 568. Solid arrowhead: false EEA1 and Rab7 co-localization which by CLEM appeared a small fold in the cryosection. Open arrowhead: closely apposed EEA1 and Rab7 staining. Corresponding CLEM and EM images on the right show an early endosome with associated vesicles. D, Another example of Rab7 + EEA1 positive area. EEA1 localizes over the early endosomal vacuole whereas Rab7 fluorescence appears to overlap with adjacent tubulo-vesicles. E, Example of EEA1-positive, Rab7 negative organelle showing a typical late endosomal morphology. Overlayed CLEM and uncolored EM images as indicated. Gold particles represent internalized BSA^5^. EE; early endosome, LE; late endosome, M; mitochondrion, PM; plasma membrane, V; vesicle. IF scale bar 2 µm, EM scale bars 200 nm.

APPL1 is recruited to membranes through interaction with Rab5^37^ as well as via its BAR domain, which binds to curved membranes^70,71^. EEA1 is also recruited to membranes by Rab5, and by the phospholipid PI(3)P. By live-cell imaging of fluorescent-tagged constructs^34^, APPL1 and EEA1 mark distinct pools of endosomes that both receive endocytic material from the plasma membrane. Cargo present in APPL1 endosomes is either sorted into a fast recycling pathway back to the plasma membrane or transferred to EEA1 endosomes, where further sorting for recycling or degradation occurs. Concomitantly, by IF of HeLa cells we confirmed previous studies^34–36^ showing that EEA1 and APPL1 spots show little overlap (**Fig**. 1F, I), while both populations co-localize with Rab5 (**Fig**. 1D, E, I). The APPL1 spots are mostly confined to a region just below the plasma membrane (**Fig**. 1E, arrows)^34^.

These IF experiments show that the selected antibodies mark distinct endo-lysosomal sub-populations. It is unknown, however, in how far these distributions coincide with morphological definitions of endo-lysosomal subtypes.

### Immuno-labeling of selected antigens on ultrathin sections yields label for IF but not EM

Tokuyasu cryosections are routinely used for immuno-EM^72,73^. The use of mild fixatives and lack of permeabilizing agents and embedding media results in a sensitive immuno-EM method, while the negative staining procedure gives optimal membrane contrast^72,73^. More recently, cryosections were proven excellent tools for IF and on-section CLEM, since they yield a high fluorescent signal-to-background ratio^66,67,74^. We incubated Hela cells with BSA coupled to 5 nm gold particles (BSA-gold^5nm^) for 3 hours to mark endocytic compartments, then fixed cells with 4% FA for 1 hour and processed them into either 250 nm thick cryosections for IF or 90 nm cryosections for immuno-gold labeling and transmission-EM (TEM) imaging (see materials and methods)^72,73^. Sections were incubated with the selected primary antibodies against Rab5, Rab7, EEA1 and APPL1 (Table 2), followed by secondary AlexaFluor-conjugated antibodies for IF or with protein A conjugated to 10 or 15 nm gold particles for immuno-EM, using our established protocols^72^.

We first tested several dilutions of the commercially available Rab5, Rab7, EEA1 and APPL1 antibodies in a single labeling approach using IF on cryosections (data not shown). Using the selected dilutions indicated in Table 2, all 4 antibodies gave a clear and distinct labeling pattern of well-defined puncta (**Fig**. 2A-C). Rab5 partially co-localized with EEA1 and APPL1 and Rab7 co-localized with CD63 (**Fig**. 2A-C). However, when using these same antibodies (in dilutions from 1:10 to 1:100) in our immuno-EM protocol, we found no significant specific labeling for any of these antigens (**Fig**. 2D, E). As positive control and to mark early and late endo-lysosomal stages we combined labelings with antibodies against Hrs (an ESCRT-0 component marking early endosomes) and CD63 (marking late endosomes and lysosomes), with proven reactivity in immuno-EM^75,76^. Since the primary antibodies for Rab5, Rab7, EEA1 and APPL1 are identical in IF and EM, the discrepancy in labeling is likely explained by the use of different reagents and dissimilar post-labeling procedures. An important conclusion from these data is that IF labeling of cryosections - instead of using whole cells as in conventional IF studies - does allow detection of these antigens. This opens the way for on-section CLEM.

### An optimal fixation protocol for fluorescence labeling and ultrastructure

In on-section CLEM methods a section is first viewed by fluorescence microscopy and then by EM. Recent developments by us^66,77^ and others^67,78,79^ have improved the accuracy and correlation efficiency of on-section CLEM to such an extent that the fluorescent signal can be directly inferred to EM sections. Contrary to IF studies, however, preservation of ultrastructure is key for interpretation of EM data. As shown in **figures** 1 and 2, the antibodies for Rab5, Rab7, EEA1 and APPL1 work well in IF of HeLa cells fixed in 4% FA for 15 or 60 minutes, respectively (**Fig**. 1 and 2). This short fixation time, however, generally results in a poorly preserved EM ultrastructure. Stronger fixatives increase ultrastructure but often abolish antigenicity. To establish the optimal balance between fluorescent signal and conservation of ultrastructure, we tested 8 different fixation regimes (**Table** 1). Fixed HeLa cells were embedded in gelatin and processed into 90 nm cryosections and either fluorescently labeled and imaged for IF, or contrasted with uranyl acetate and examined by TEM. To measure the signal in fluorescence microscopy we calculated the signal-to-noise-ratio (SNR) as mean intensity value of the 0.5% brightest pixels divided by the mean intensity value of the reverse selection. To classify EM morphology, a panel of lab members blindly ranked the resulting images based on the cohesion of the cytoplasm and the visibility and sharpness of membranes.

We found that a 30-minute fixation with 4% FA yielded the best signal-to-noise ratio in fluorescence microscopy, however, preservation of EM morphology was very poor in these conditions (Table 1, **Fig**. S1). Adding 0.2% glutaraldehyde (GA) greatly improved EM morphology (Table 1, **Fig**. S2), but deteriorated the fluorescence signal even when we quenched the GA autofluorescence with NaBH_4_ before labeling. Using only FA fixation, we found that increased fixation times very rapidly decreased the fluorescence microscopy signal for all antibodies (Table 1), with a notable decline already after 1 hour. As optimal compromise between fluorescence signal and morphology, we selected a mild fixation of 4% FA for 1 hour as best fixative for CLEM.

### CLEM of Rab5 and Rab7 reveals differential distributions over early to late endo-lysosomal compartments

We then executed a full CLEM experiment by performing IF and EM on the same section (**Fig**. 3A). We incubated HeLa cells for 3 hours with BSA-gold^5nm^, fixed cells for 1 hour in 4% FA and then immediately scraped cells to prepare roughly 1 mm^3^ gelatin blocks that were plunge-frozen and stored in liquid nitrogen. We collected 90 nm cryosections from these blocks on an EM carrier grid and labeled these with the selected primary antibodies, AlexaFluor-tagged fluorescent secondary antibodies and DAPI to stain the nuclei. The fluorescently labeled sections were imaged in a widefield fluorescence microscope and tilesets of the ribbon of sections present on the grid were collected^66,67^. After imaging, the grids were stained with 2% uranyl acetate, which is our normal contrasting procedure for immuno-EM. In the EM, we again collected large image tilesets at 43,000x magnification to resolve endo-lysosomal membranes in great detail and stitched these together using Etomo post-processing software. The nuclei, in fluorescence microscopy identified by DAPI signal and in EM by morphology, served as numerous unique reference points which made the correlation between the IF and EM images highly accurate (**Fig**. S3). Combining these large datasets of fluorescence microscopy and EM allows correlation of hundreds of single organelles from tens of different cells in one sample^66^, which greatly increases the throughput of on-section CLEM.

We first applied high-throughput CLEM on sections of HeLa cells double-labeled for endogenous Rab5 and Rab7 (**Fig**. 3). After performing correlation, we could readily define Rab5, Rab7 and Rab5/7-positive puncta by their succinct fluorescent signal (**Fig**. 3B). We then correlated these spots to EM ultrastructure and classified the underlying structures as vesicle, early or late endosome or lysosome. The precise morphological criteria of these distinct endo-lysosomal intermediates are based on a wealth of previous EM and immuno-EM studies from many distinct laboratories^14,17,20,41,42,47,49,54,80–82^ and summarized in the materials and methods section. Based on these ultrastructural definitions, our on-section CLEM approach localized Rab5 mainly to vesicles and tubules (70%) and early endosomes (19%). The Rab5-positive vesicles-tubules were 100-200 nm in diameter and often found near the plasma membrane. They occasionally contained internalized BSA-gold^5nm^ and in 23% of the cases displayed a characteristic clathrin coat, by which they meet all the criteria of endocytic vesicles. Rab5-positive early endosomes were in 43% of the cases positive for BSA-gold^5nm^ and in 57% the characteristic flat, bi-layered clathrin coat was visible within the plane of sectioning (**Fig**. 3C). Our results thereby match existing literature, where Rab5 has been described both on endosomal vacuoles^65^, as well as on endocytic vesicles^83^. In addition, we found a small fraction of Rab5 staining over compartments that meet the criteria of late endosomes (**Fig**. 3D).

Rab7 showed a very different distribution pattern and was mainly associated with late endosomes (33%) and lysosomes (39%, **Fig**. 3D). During the 3 hours of uptake, 86% and 44% of Rab7-positive late endosomes and lysosomes, respectively, was reached by BSA-gold^5nm^. These numbers reflect the kinetics by which these distinct stages of the endocytic pathway are reached. They can, however, not be taken as absolute numbers, since some negative organelles may contain BSA-gold^5nm^ outside the plain of sectioning. The Rab7 distribution pattern is in line with previous studies^14,63^ and supports its role as an organizer of the late endocytic pathway. Only 6% of all analyzed compartments was positive for both Rab5 and Rab7, the organelles underlying these double-labeled puncta had both early and late endosomal characteristics.

By correlating a large number of organelles (n = 101) we systematically categorized the distribution patterns of Rab5 and Rab7 (**Fig**. 3D). This quantitative analysis showed that endogenous Rab5 and Rab7 are reversely distributed over early and late endocytic compartments. Overall, these distributions correspond to the known functions and localizations of these Rabs and thereby validate the feasibility of our approach. Of note, we show that in these standard conditions in HeLa cells the majority of Rab5 is present on small endocytic vesicles rather than on early endosomal vacuoles, which are two functionally distinct stages of early endocytosis.

### CLEM localizes EEA1 and APPL1 to morphologically different compartments

Next, we performed on-section CLEM on HeLa cells double-labeled for APPL1 and EEA1 (**Fig**. 4). Using the same morphological definitions as for Rab5 and Rab7, we found that APPL1 and EEA1 show very different localization patterns and seldomly overlap (circa 5% of organelles, **Fig**. 4A, D). APPL1 staining consistently marked tubulo-vesicular membranes of 100-150 nm diameter, sometimes forming a cluster together (**Fig**. 4B). This is consistent with a previous pre-embedding immunolabeling study^34^, showing that APPL1 was present on vesicular structures rather than ‘classical’ early endosomes consisting of a vacuole and associated tubules. Consistent with literature we will refer to these APPL1 vesicles as APPL1 endosomes^34,35^. We found APPL1 endosomes often in the vicinity of the plasma membrane, which matches the pattern observed in IF. Analysis of 53 APPL1-labeled organelles, showed that 30% of the APPL1 endosomes exhibited a clathrin coat and 25% contained internalized BSA-gold^5nm^, confirming their endocytic nature. Since accumulation of APPL1 on endosomes coincides with loss of clathrin^36^, the APPL1 endosomes that still display a clathrin coat are likely freshly derived from the plasma membrane, while those without are older. Notably, except for clathrin coats, APPL1 endosomes lacked any morphological characteristics; they formed no membrane buds or branches and their lumen lacked any internal membranes. EEA1, on the other hand, was found on a variety of endosomal organelles, ranging from small (100 – 200 nm) endocytic vesicles to classical early endosomes containing ILVs and, unexpectedly, to late endosomes and lysosomes with internal membranes and dense content (**Fig**. 4B). Quantification showed that about half of the fluorescent EEA1 puncta was localized to the later stages of the endo-lysosomal system (**Fig**. 4C). In case co-localization of APPL1 and EEA1 was seen by IF, this often revealed an early endosome with a vesicle in close vicinity, representing separate EEA1 and APPL1 labeled compartments, respectively (**Fig**. 4D).

### EEA1 localizes to late endosomal compartments that lack Rab5

EEA1 is generally considered as an early endosomal protein, since it binds Rab5^84^ and interacts through its FYVE domain with PI(3)P^85,86^, which is enriched on early endosomes^87^. The relatively high percentage of EEA1 on late endosomal compartments (**Fig**. 4C) was therefore unexpected, also since we found Rab5 mostly on endocytic vesicles and early endosomes, not on late endosomes (**Fig**. 3D). Collectively, these data suggest that EEA1 is present on Rab5-positive early endosomes and Rab5-negative late endosomal compartments. To address this in more detail, we performed a CLEM double-labeling for EEA1 and Rab5 (**Fig**. 5A) and specifically quantitated their co-localization per category of endosomal organelle: 100-200 nm vesicles-tubules, early endosomes and late endosomes (**Fig**. 5B). Since Rab5 labeling on lysosomes is negligible, we did not take these along. Of the analyzed 100-200 nm vesicles-tubules, 52% exclusively displayed Rab5, 21% was positive for EEA1 and 27% displayed both markers (**Fig**. 5B). Likely, part of the Rab5-only vesicles will represent APPL1 endosomes. Most of the bona-fide early endosomes, i.e. with a distinctive vacuolar part and containing ILVs, were positive for EEA1 (51%) or both Rab5 and EEA1 (44%). The vast majority of late endosomes was EEA1-positive (77%), whereas 18% was positive for both markers. On both early and late endosome, only 5% was positive for Rab5 only. These data clearly show that Rab5 has a distribution distinct from EEA1: Rab5 is mostly confined to early-stage endocytic vesicles and early endosomes, whereas EEA1 also labels late endosomal compartments. Possibly, EEA1 remains on late endosomes through its interaction with PI(3)P^8,12^ after Rab5 has dissociated.

### EEA1-positive late endosomal compartments can be surrounded by Rab7 vesicles

Since the dissociation of Rab5 coincides with the recruitment of Rab7^12,13^, we next investigated if the EEA1-positive/Rab5-negative late endosomes (**Fig**. 5B) contain Rab7. IF of double-labeled, 90 nm thin cryosections of HeLa cells revealed minimal Rab7 and EEA1 co-localization, in accordance with previously reported IF on whole cells^88^. Less than 5% of the EEA1 spots overlapped with Rab7 (**Fig**. 5C), part of which by CLEM appeared as false-positive, i.e. fluorescence caused by a fold in the section or dirt particle (**Fig**. 5C, solid arrowhead). When EEA1 and Rab7 fluorescent spots were found adjacent rather than overlapping (**Fig**. 5C, open arrowhead), CLEM analysis revealed an early or late endosome positive for EEA1 with the Rab7 signal correlating to associated tubulo-vesicular structures (**Fig**. 5C, D). These endosome-associated vesicles and tubules possibly represent Rab7/Retromer-positive recycling tubules, emerging from endosomes where EEA1 is still maintained by the lipid PI(3)P^28,89^. We readily found EEA1-positive late endosomes without Rab7 (**Fig**. 5E). Combined, these data suggest the existence of a pool of EEA1-positive endosomes that is not labeled by Rab5 or Rab7, but may be forming Rab7-positive recycling membranes.

### EEA1 localization to late endosomes is conserved between cell lines

Given the unexpected association of EEA1 with late endosomes, we wanted to investigate whether this is a specific feature of Hela cells or is representative for the general distribution of EEA1. To address this question, we performed on-section CLEM of EEA1 in HEPG2 (human hepatoma), A549 (human adenocarcinoma from alveolar basal epithelium) and HT1080 (human fibrosarcoma) cell lines (**Fig**. 6A-C). For comparison, we combined it again with APPL1 staining. In all cell lines, we found EEA1 on early as well as late endosomal compartments (**Fig**. 6D). The relative distribution differed between the cell lines, but in all the presence of EEA1 label on late endosomes or lysosomes was substantial, specifically 45%, 29% and 15% in HEPG2, HT1080 and A549 cells, respectively. Furthermore, in contrast to HeLa cells, in A549 and HT1080 cells, a considerable portion of EEA1 (44% and 41%, respectively) was found on 100-200 nm vesicles, presumably of endocytic origin. Similar to Hela cells, APPL1 in all cell lines showed a consistent localization to tubulo-vesicular membranes with limited overlap with EEA1 (**Fig**. 6D).

These quantitative and ultrastructural data show that EEA1 and APPL1 label structurally very different compartments and that EEA1 has a more widespread distribution than thus far anticipated. Additionally, the data show that the relative distribution of endosomal “marker” proteins over distinct endosomal compartments can differ between cell lineages.

**Figure 6.**
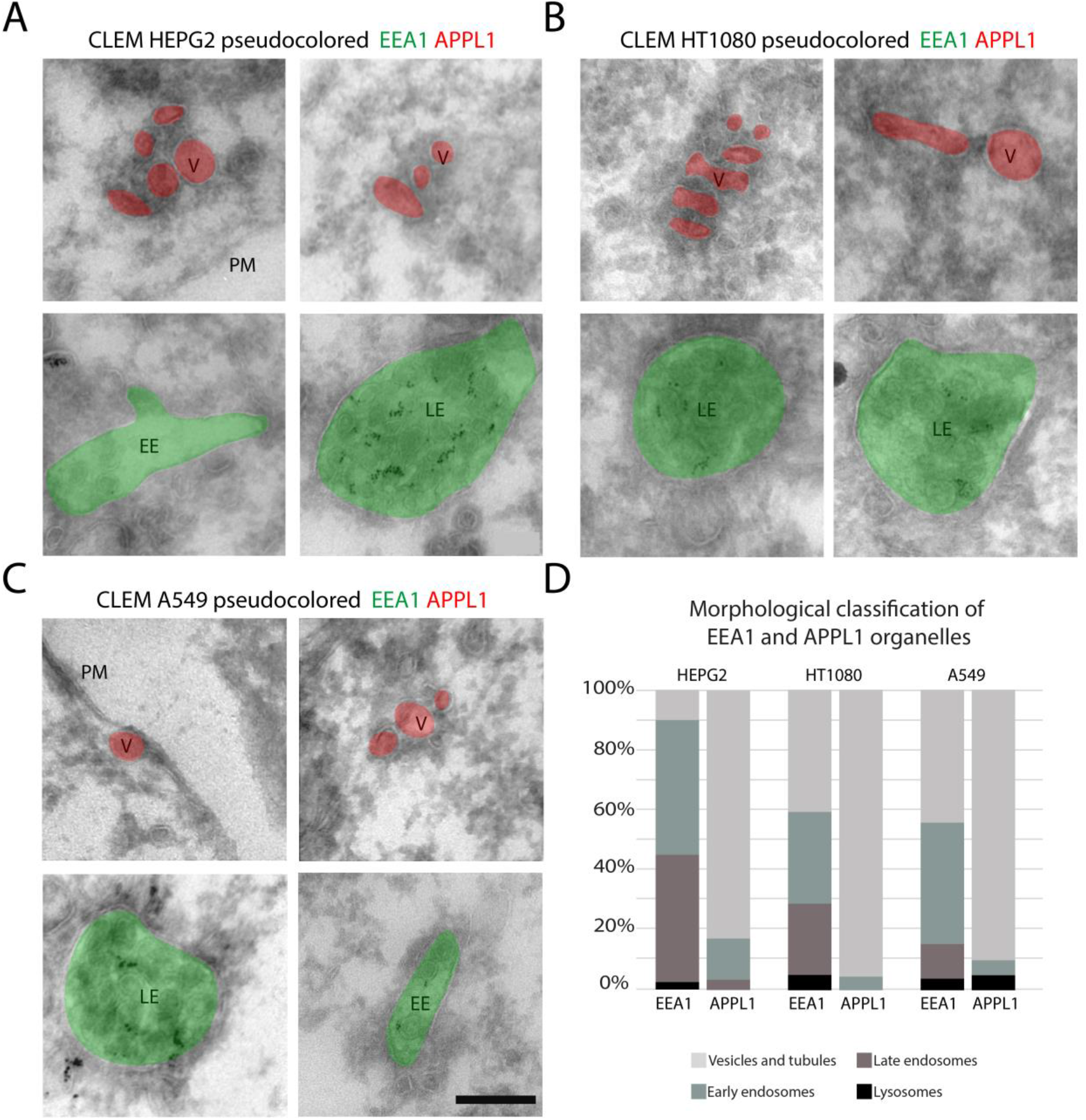
Organelle specific distributions of EEA1 and APPL1 are conserved between cell types. Indicated cell lines were prepared for CLEM as in Figure 3. A-C: Pseudocolored EM images based on double IF microscopy of EEA1 and APPL1. Original images are shown in Fig. S6. Gold particles represent internalized BSA^5^. D: Relative distribution of EEA1 and APPL1 over distinct endo-lysosomal compartments in indicated cell lines. APPL1 is consistently found over small vesicles. EEA1 is consistently found on late endosomes albeit with distinct relative distributions. HEPG2 n= 40 and 29, HT1080 n = 59 and 44, A549 n = 52 and 40 organelles for EEA1 and APPL1, respectively. EE; early endosome, LE; late endosome, PM; plasma membrane, V; vesicle. Scale bars 200 nm.

## Discussion

In this paper we introduce a high throughput CLEM method to quantitatively study the subcellular localization of selected proteins. By correlating hundreds of spots we show the quantitative localization of different combinations of proteins, with 65-120 nm correlation accuracy and transmission EM resolution. Moreover, by applying this technology to proteins that cannot be detected by conventional immuno-EM, we unleash the possibility for high resolution localization for an entirely new set of proteins. We make use of ultrathin cryosections which are traditionally used for immuno-EM but are also highly compatible with IF imaging. This makes it possible to label proteins on-section for IF imaging and subsequently overlay this signal to EM images of the same section. High throughput data are obtained by making stitched high magnification images of sections in IF and EM, which contain numerous reference points (i.e. nuclei) for quick and accurate alignment of the datasets (**Fig**. S3, ^66^). We demonstrate the power of our approach by revealing the sub-cellular distributions of key proteins of the endo-lysosomal system, i.e. Rab5, EEA1, APPL1 and Rab7. Our data reveal novel information on the spatial distribution of these proteins over distinct endo-lysosomal compartments, revealing new insights in the composition of the endo-lysosomal system and with consequences for their use as markers for specific endo-lysosomal compartments.

Our data clearly demonstrate that proteins detected by IF cannot always be localized by immuno-EM (**Fig**. 2). Since the primary antibodies used in these experiments were identical, this discrepancy is likely due to the use of different labels and/or differences in sample preparation between fluorescence microscopy and EM methods. Indeed, colloidal gold particles show limited penetration into cryosections^90,91^, which would explain a lower signal-to-noise ratio than obtained with fluorescent probes penetrating the entire section. Alternatively, or additionally, the post-labeling approach for EM may result in some loss of antibody-gold complexes, for example during washes in H_2_O or post-fixation with uranyl. Labeling for immuno-EM can be enhanced by pre-embedding labeling, silver enhancement or peroxidase stains, but these methods affect or obscure morphology, are not quantitative and limit the number of specific proteins that can be labeled simultaneously. The here presented on-section CLEM method therefore offers a strong and attractive alternative to detect proteins that fail to label in immuno-EM.

The first step towards CLEM is to overcome the distinct requirements for fluorescence microscopy (optimized for a high labeling signal) and EM imaging (optimized for high ultrastructural preservation)^49,92^. Testing different fixation conditions, we found a striking effect of FA fixation duration on IF labeling (**Table** 1). Fixation times longer than 1 hour significantly decreased the signal for all 4 proteins under study, i.e. Rab5, EEA1, APPL1 and Rab7. As optimal compromise between fluorescence signal and morphology, we selected a mild fixation of 4% FA for 1 hour as best fixative for CLEM. Although in these conditions EM ultrastructure is not maximally preserved, all defining characteristic features of endo-lysosomal compartments are readily visible, allowing an accurate identification based on their morphology. We suggest to test these conditions for each antigen to be studied by CLEM, by which it is recommended to seek for the strongest fixation possible without significant loss of signal.

Our studies reveal insightful new information on the localization of the proteins under study. Double-labeling of Rab5 and Rab7 showed a complementary distribution over early and late endosomal compartments, respectively, with very limited overlap. Notably, in the HeLa cells we used for the majority of our experiments the vast majority of Rab5 (70%) was associated with small, 100 – 200 nm singular endocytic vesicles and tubules rather than with early (19%) endosomes. The relative distribution of Rab5 over these distinct compartments may vary between cells and conceivably between different experimental conditions. How the IF pattern in different conditions translates to these distinct compartments is important for the interpretation of experiments, since endocytic vesicles and early endosomes are functionally distinct compartments, with early endosomes being complex structures with different molecular and functional membrane domains that enable cargo sorting^9,41,59,83^. Rab5 is now commonly referred to as early endosome marker. Based on our data ‘marker for early endocytic compartments’ would be more correct. Rab7 was validated as suitable and specific marker for late endocytic compartments, which encompassed both late endosomes as well as lysosomes (**Fig**. 3D, **Fig**. 7). Occasionally we observed Rab7-positive vesicles close to or associated with an endosomal vacuole (**Fig**. 5C). These likely represent Rab7/Retromer-positive recycling tubules emanating from endosomal vacuoles^14^.

**Figure 7.**
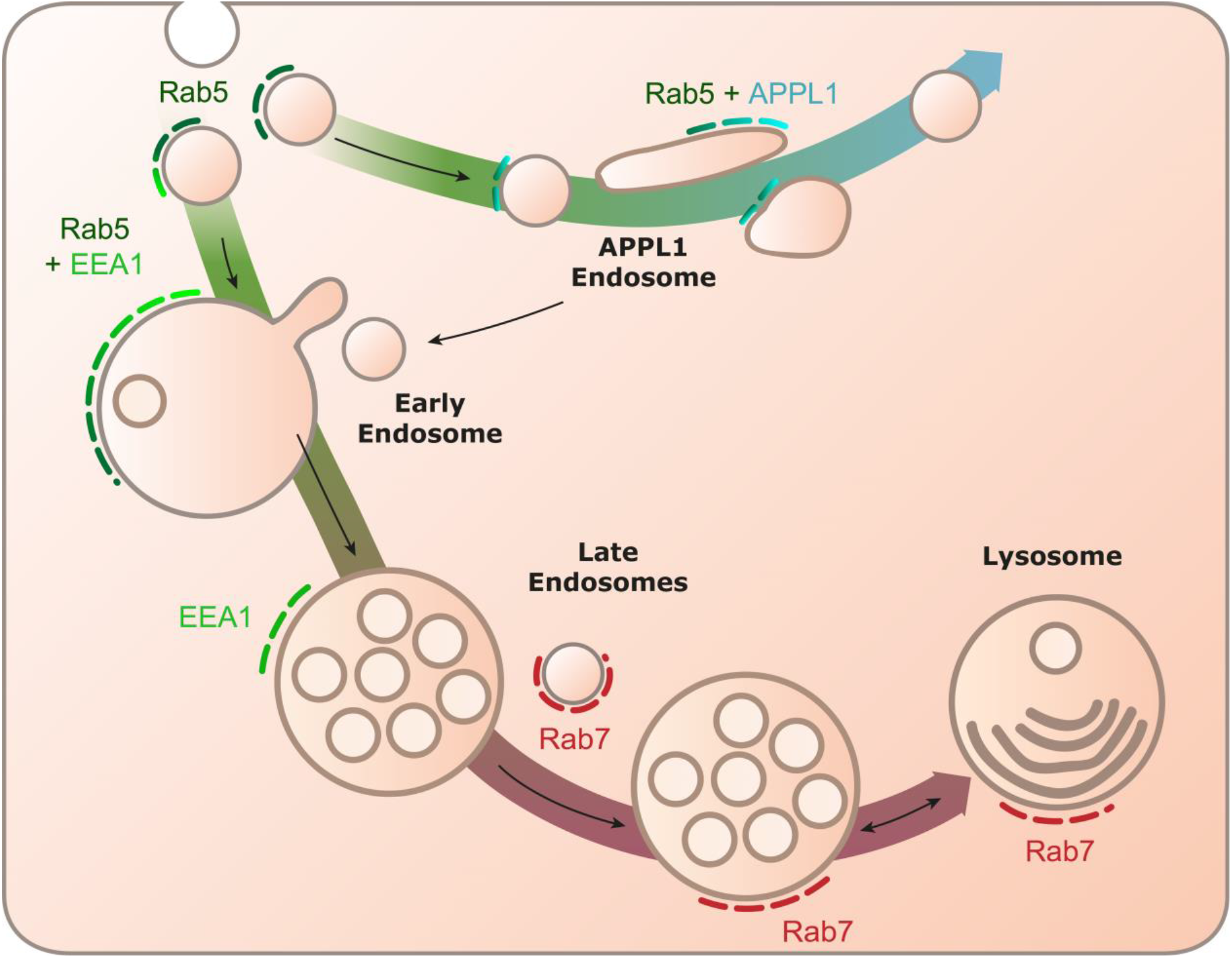
Schematic organization of the endo-lysosomal system based on CLEM localization of Rab5, Rab7, EEA1 and APPL1. Our data show that Rab5 is mostly present on endocytic vesicles and early endosomes. Rab5 and EEA1 co-localize on early endosomes, whereas EEA1 additionally resides on late endosomes. Rab5 overlaps with APPL1 l on an entirely distinct population of endosomes consisting of clusters of small tubulo-vesicles. Late endosomes and lysosomes are marked by Rab7. Large arrows signify recycling (upper) and degradative (lower) pathways, small arrows indicate maturation, vesicle trafficking or fusion events.

Focusing on the Rab5 effectors APPL1 and EEA1, we confirmed by IF that these two proteins mark separate pools of endosomal organelles^34^, and by EM show that these pools represent morphologically distinct membranes (**Fig**. 4C, **Fig**. 7). APPL1 is consistently found on small vesicles (APPL1 endosomes) that are mostly oval shaped, 100-150 nm in diameter and length, and sometimes clustered together. Apart from an occasional clathrin coat these endosomes had no distinguishing features like tubules or ILVs, nor did they contain any discernable content, except when cells were incubated with BSA-gold^5nm^ before fixation. The consistent association of APPL1 with small, high curvature vesicles and tubules could be explained by its BAR domain, which promotes membrane curvature^71,93,94^. By both IF and EM (**Fig**. 1C and **Fig**. 4C) we typically found the APPL1-positive vesicles close to the plasma membrane. APPL1 has been proposed to serve as adaptor for membrane receptors and signaling proteins^37^. How the small APPL1 endosomes regulate cargo sorting and recycling^34,37^ needs to be established. Only a very small fraction (5%) of APPL1 localized with EEA1 by CLEM. These spots by EM often appeared as a typical, small APPL1 endosome close to an EEA1-positive endosomal vacuole. EEA1 showed a much wider distribution than APPL1, ranging from 100-150 nm endocytic vesicles to early and late endosomes and even lysosomes (**Fig**. 4D).

A striking finding was that a significant portion of EEA1 localizes to late endocytic compartments. We found this in Hela cells as well as HEPG2, A549 and HT1080 cell lines. In all cell lines, EEA1 was present on early as well as late endosomal compartments, albeit with distinct relative distributions. Hence, the data obtained from IF cannot simply be extrapolated to a functionally defined compartment, like an early or late endosome, but differs between cells and most likely also between experimental conditions. Furthermore, these data necessitate reassessment of the widespread use of EEA1 as marker for early endosomes: Our data show that EEA1 is an appropriate marker for early and late endosomes. Again, this is an important finding with respect to the interpretation of co-localization data in IF. Keeping in mind that EEA1 is present on early and late endosomes may completely change the interpretation and impact of such data. In studies that require specific detection of early endosomes we recommend double-labeling between EEA1 and Rab5, since the combination of these two markers more specifically labels early endosomes (**Fig**. 5A, B).

Since EEA1 is recruited to membranes by both Rab5 and PI(3)P, it is possible that presence of EEA1 is maintained on maturing endosomes by PI(3)P. PI(3)P is known to be present on maturing endosomes, until it is phosphorylated to PI(3,5)P2^8,12^. Furthermore, we occasionally find organelles positive for both EEA1 and Rab7. These endosomes often display recycling tubules and vesicles. This suggests that EEA1 persists on maturing endosomes up to Rab7-Retromer based recycling, independent of Rab5. Additionally, we found a population of EEA1-positive endosomes not labeled by either Rab5 or Rab7. Endogenous tagging approaches to label simultaneously for Rab5, Rab7 and EEA1 are ideal to further study this population, and volumetric EM methods can be employed to exclude that Rab5 or Rab7 subdomains are missed on organelles in thin sections^54,65^. Additionally, further mechanistic research can reveal whether EEA1 still actively tethers incoming Rab5 vesicles to late endosomes.

Concluding, we present a high throughput CLEM application as attractive and quantitative method to localize proteins in a morphological context when classical EM labeling schemes fail. Applications of this method are numerous, since it can in principle correlate any IF signal visible in sections, either obtained through immuno-labeling or otherwise. Furthermore, our method can be combined with other imaging methods^95^, including super-resolution microscopy, since cryosections are compatible with Photo-Activatable Light Microscopy (PALM) or Stimulated Emission Depletion (STED) imaging, provided suitable fluorophores are used^96^. To increase the throughput of future studies automatized computational correlation using fiducial markers^66^ is an option, as is machine learning-based feature detection and automatic cross-correlation^97^. In any way, high throughput on-section CLEM can connect molecular composition to morphology and provide novel understanding without the need for new equipment or reagents beyond those required for conventional immuno-EM and IF.

## Materials and methods

### Antibodies

### Cell culture

HeLa, A549, HepG2 and HT1080 cells were cultured in Corning T-75 cell culture flasks placed in a 5% CO_2_ incubator at 37°C. Cells were grown in Dulbecco’s Modified Eagle’s Medium (DMEM) supplemented with 10% fetal calf serum, 100 U/ml Penicillin and 100 ug/ml Streptomycin. For IF, cells were seeded on 15mm coverslips in a 24-well plate. For EM and CLEM samples, cells were grown in 6cm culture dishes and incubated with BSA-gold^5nm^ particles (Cell Microscopy Core, UMC Utrecht) in full DMEM for 3 hours prior to fixation.

### Immuno-fluorescence

Cells on coverslips were fixed with 4% FA for 15 minutes followed by 3 PBS washes and permeabilization in TritonX-100 0.1% in PBS for 10 minutes. Blocking was performed in 1% BSA in PBS for 10 minutes before incubation with primary antibodies in 1% BSA for 1 hour at room temperature. Coverslips were incubated with secondary antibodies for 30 minutes at room temperature and mounted in Prolong Diamond with DAPI.

Samples were imaged on a LSM700 Leica confocal microscope using a 63x oil objective. Images were recorded as single slices with pinhole size at 1 airy unit for each channel. Images were analyzed in FIJI using the ComDet 5.5 plugin (Eugene Katrukha, Cell Biology, Utrecht University) and a custom macro. For the analysis resulting in figure 1G, 124 cells from 2 independent replicates were analyzed, 102±71 and 391±218 mean±SD particles were found per cell for Rab7 and Rab5, respectively. The percentage of co-localized particles was calculated per cell. For averages and standard deviations, see Table S1. For figure 1H, 147 cells from 2 independent replicates were analyzed, 258±154, 119±84 and 106±42 particles were found per cell for Rab7, CD63 and Cathepsin D, respectively. For figure 1I, 352 cells from 2 independent replicates were analyzed, 121±60, 211±129 and 344±206 particles were found per cell for EEA1, APPL1 and Rab5, respectively.

### Sample embedding

For CLEM and Tokuyasu immuno-EM, sample preparation and sectioning were performed as previously described^72,73^. A detailed protocol is available in supplemental Note S1. In short, cells were fixed by adding 4%FA in 0.1M Phosphate Buffer (PB) 1:1 to culture medium to reduce osmotic shock. After 5 minutes, medium and fixative were replaced with 4% FA in 0.1M PB for 1 hour at room temperature unless otherwise indicated. Fixative was washed off and quenched with PBS + 0.15% glycine. Cells were detached from the culture dishes using scrapers and collected in PBS with 1% Gelatin. After pelleting cells, 1% gelatin was replaced by 12% gelatin at 37°C and cells were pelleted again. The pellets were solidified on ice, cut into smaller blocks and infused with 2.3M sucrose overnight at 4°C. The smaller blocks were mounted on pins and stored in liquid nitrogen.

The gelatin-embedded cells were cryosectioned to 90 nm thick sections at -100°C on a DiATOME diamond knife in a Leica ultracut cryomicrotome. Sections were picked up and deposited on formvar- and carbon-coated grids using 2.3M sucrose and 1.8% methylcellulose mixed 1:1.

### Immuno-EM

We performed the immunolabeling procedure as developed in our lab and described in detail in^72^. Sections on grids were washed using PBS at 37°C for about 30 minutes to remove the 2.3M sucrose and 1.8% methylcellulose mixture. After washing and blocking steps, we performed labeling using primary antibodies, followed by an incubation with bridging antibodies where needed. Grids where then incubated with Protein A conjugated to gold particles of 10 or 15 nm size (Cell Microscopy Core, UMC Utrecht). Grids were postfixed for 5 minutes using 2% Uranyl acetate pH 7.0, followed by Uranyl acetate/methylcellulose mixture, pH 4.0 for 10 minutes at 4°C.

Imaging was performed on a Tecnai T12 TEM using serialEM software.

### On-section CLEM procedure

A step-by-step protocol of the CLEM procedure, including reagents and footnotes is included as a supplemental file to this manuscript (Supplementary Note S1). In short; sections on grids were washed using PBS at 37°C for 30 minutes, followed by short PBS washes, a blocking step and incubation with primary antibodies for 1 hour at room temperature as described in the previous paragraph. Sections were then incubated with fluorescent secondary antibodies, as well as DAPI for 30 to 90 minutes. After 30 minutes, 2-3 grids were washed in PBS 5 times, submersed in 50% glycerol and sandwiched between a clean coverslip and slide glass^103^, sections facing the coverslip. These grids were then imaged on a Leica Thunder fluorescence microscope using a 100x oil objective. Stitched images were collected, providing a complete view of all sections on a grid. The grids were retrieved by removing the oil from the coverslip and gently dislodging the coverslip from the slide glass. The grids were washed in PBS and the conventional immuno-EM protocol was resumed with incubation in 1% GA for 5 minutes, washes in H_2_0 and postfixation using uranyl acetate. We processed only 2-3 grids at a time from secondary incubation onward to reduce time on 50% glycerol and diffusion of labeling.

After sample preparation for EM, sections are imaged in a Tecnai T12 TEM using serialEM^79^ software. We selected the regions for EM tileset images based on the fluorescence images of the same section. After acquisition of the EM images, the data was transferred to a workstation computer and stitched together using Etomo montage blending software^104^. The stitched EM image tileset and the corresponding fluorescence image were loaded into Adobe photoshop 2019 and aligned based on DAPI signal and morphology of nuclei. Images were linearly resized, rotated or moved in x and y to achieve best visual overlay. We also performed landmark-based correlation using the ec-CLEM plugin^78^ in Icy software^105^ to assess the accuracy of our overlays (**Fig**. S3). The landmarks were again based on DAPI and nuclear morphology, and yielded a predicted error of 60-120 nm across the correlated image (**Fig**. S3E). Correlated images were exported as .tif files and loaded into FIJI^106^ to select and individually crop organelles. These were categorized based on morphological criteria, resulting in organelle distributions.

### Definition of endo-lysosomes by EM morphology

A wealth of EM images collected over the last decades, by many different labs and by many different methods, has resulted in general morphological criteria of endo-lysosomal compartments, see for some examples from literature^20,41,44,54^. Based on these collective studies we here defined early endosomes as irregularly shaped electron-lucent vacuoles containing <6 intraluminal vesicles (ILVs), often displaying a flat clathrin coat (which contains Hrs) and/or associated tubules; Late endosomes as globular shaped vacuoles with a relatively electron dense content and containing >6 ILVs; Lysosomes as irregularly shaped and sized vacuoles with a variable, mixed content of ILVs, amorphous, electron-dense material and degraded membranes that can form onion-like concentric rings^54^. Structures smaller than 200 nm in diameter were designated as ‘tubulo-vesicular’, since a round profile might represent a cross-section of an elongated tubule. Although these categorizations are not absolute, they maximally represent our current knowledge on structure-function relationships and offer an objective tool to compare the distribution of the different endosomal markers using the same criteria.

## Acknowledgements

We thank our colleagues of the Cell Biology section and Centre for Molecular Medicine for fruitful discussions and feedback. We thank C. Rabouille for constructive and valuable discussion of the work. N.L. is supported by a ZonMW TOP grant [40-00812-98-16006 to J.K]. The electron microscopy within this work is part of the research program National Roadmap for Large-Scale Research Infrastructure (NEMI), project number 184.034.014, which is financed by the Dutch Research Council (NWO).

## Supplemental files

**Figure S1.**
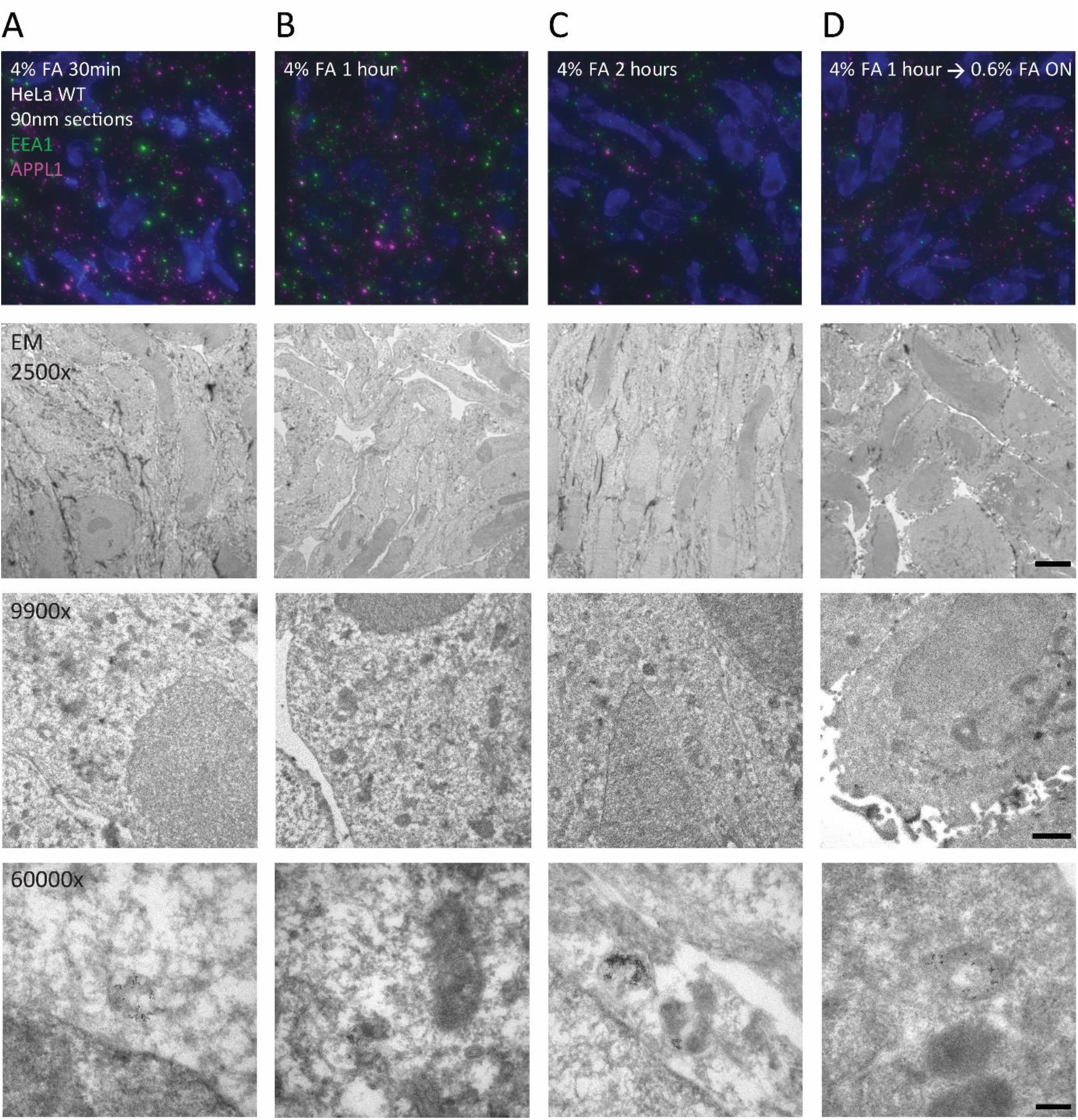
Effect of prolonged FA fixation on IF signal and EM morphology. Widefield microscopy (upper row) and EM images of 90 nm thick cryosections prepared from HeLa cells fixed according indicated protocols. Sections were fluorescently labeled for EEA1 and APPL1 (upper row) or directly prepared for EM. More than 1 hour FA fixation significantly reduces fluorescent signal. IF images are presented with identical intensity threshold settings. See Table 1 for quantifications of fluorescence microscopy signal-to-noise ratio and EM morphology. 2500x scale bar 5 µm, 9900x scale bar 1 µm, 60000x scale bar 200nm.

**Figure S2.**
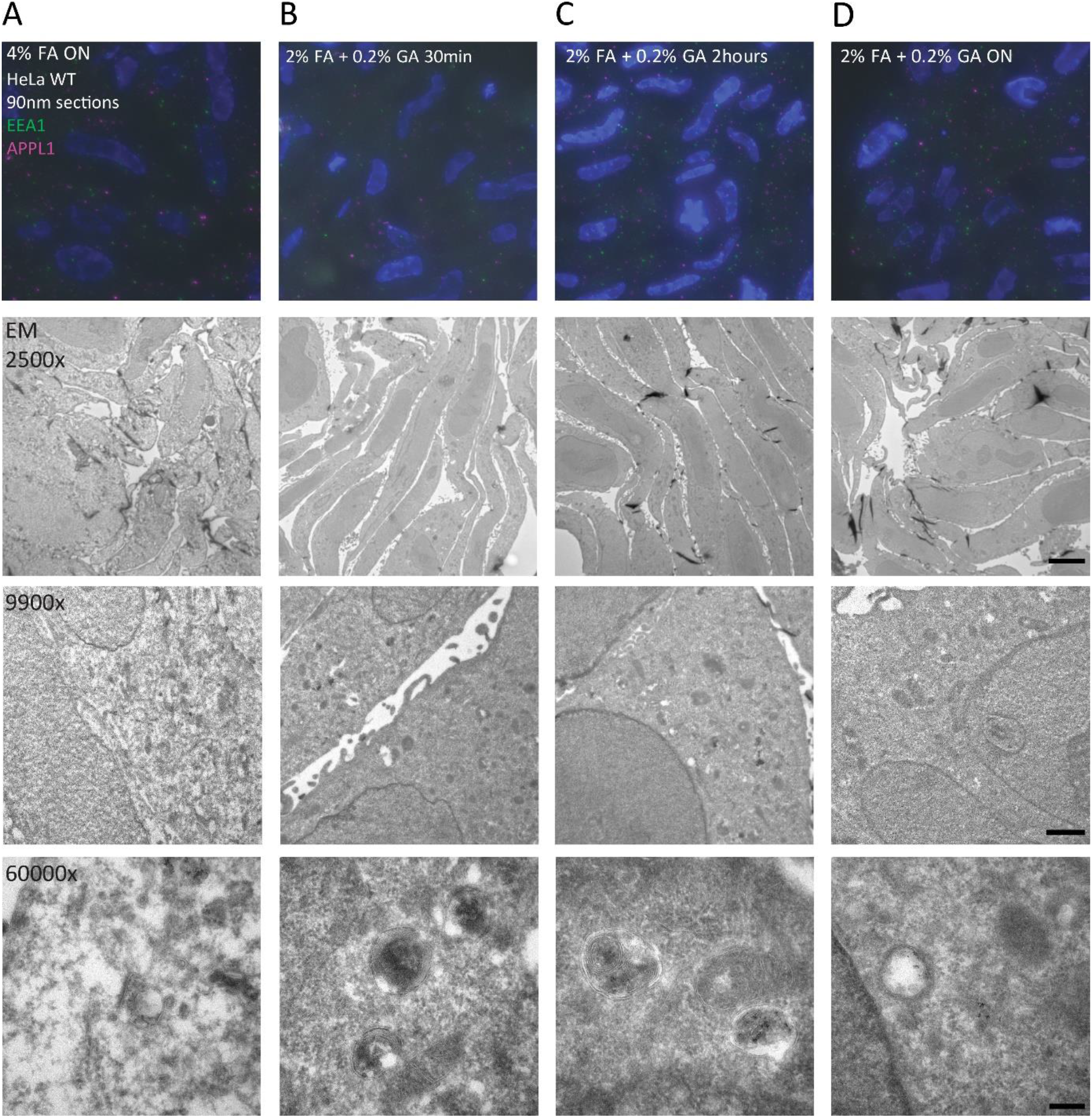
Effect of GA fixation on IF signal and EM morphology. Widefield microscopy and EM images of 90 nm thick cryosections prepared from HeLa cells fixed according indicated protocols. Sections were fluorescently labeled for EEA1 and APPL1 or directly prepared for EM. GA fixation greatly improves EM morphology, but averts the immuno-fluorescent signal. IF images are presented with identical intensity threshold settings. See Table 1 for quantifications of fluorescence microscopy signal-to-noise ratio and EM morphology. 2500x scale bar 5 µm, 9900x scale bar 1 µm, 60000x scale bar 200nm.

**Figure S3.**
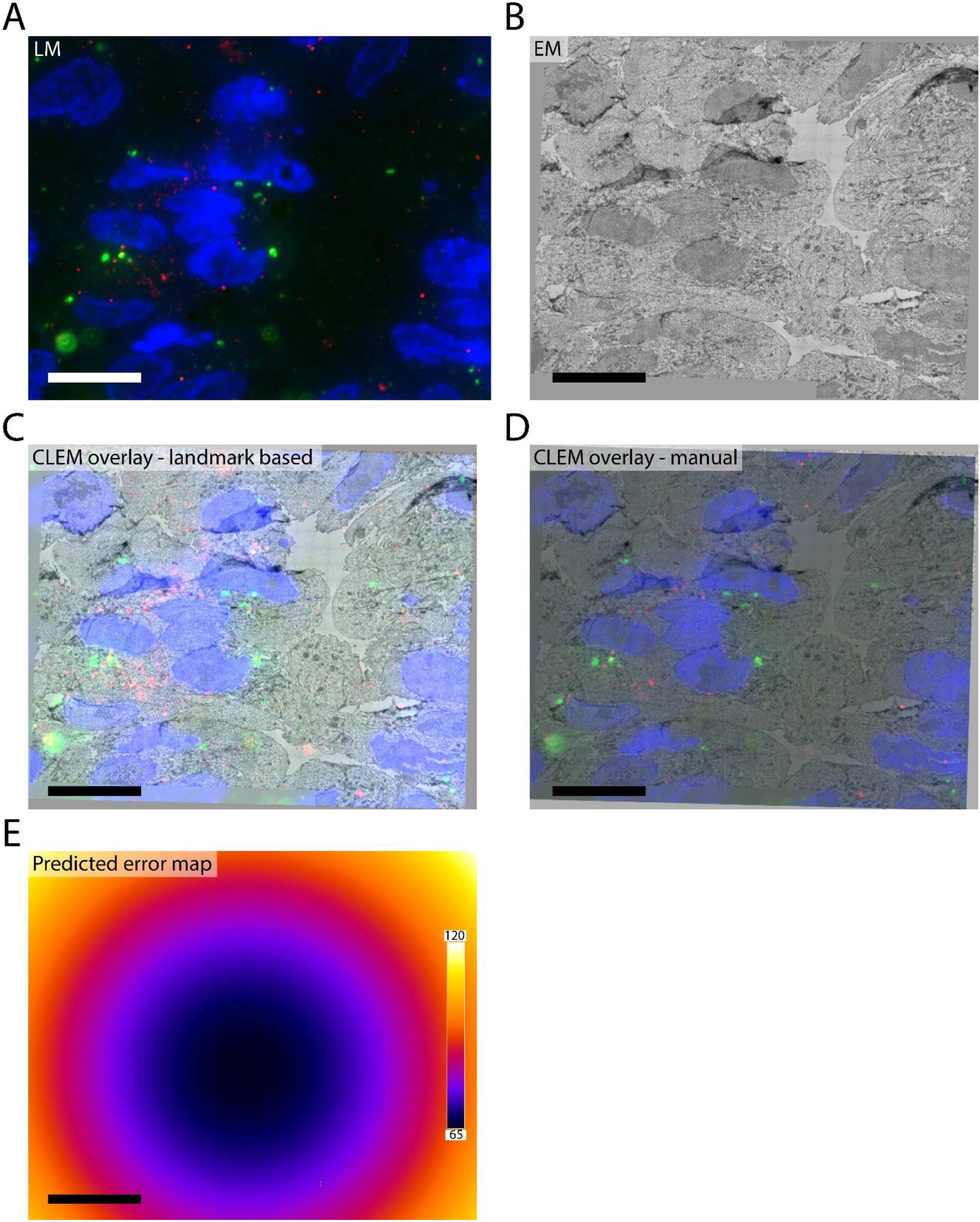
Correlation accuracy of IF to EM overlays. A, B: IF (A) and EM (B) of the same area of HeLa 90nm cryosections labeled for EEA1 (green) and APPL1 (red). C: Overlay of IF and EM based on landmarks selected on DAPI and nuclear morphology using the ec-CLEM plugin in Icy. D: Overlay based on best visual match of DAPI and nuclear morphology. Both methods yield very similar results. E: Predicted error map of overlay from C. The center of the image is accurately overlayed with a 65 nm error margin, the edges with 120nm error margin. Scale bars 10 µm.

**Figure S4.**
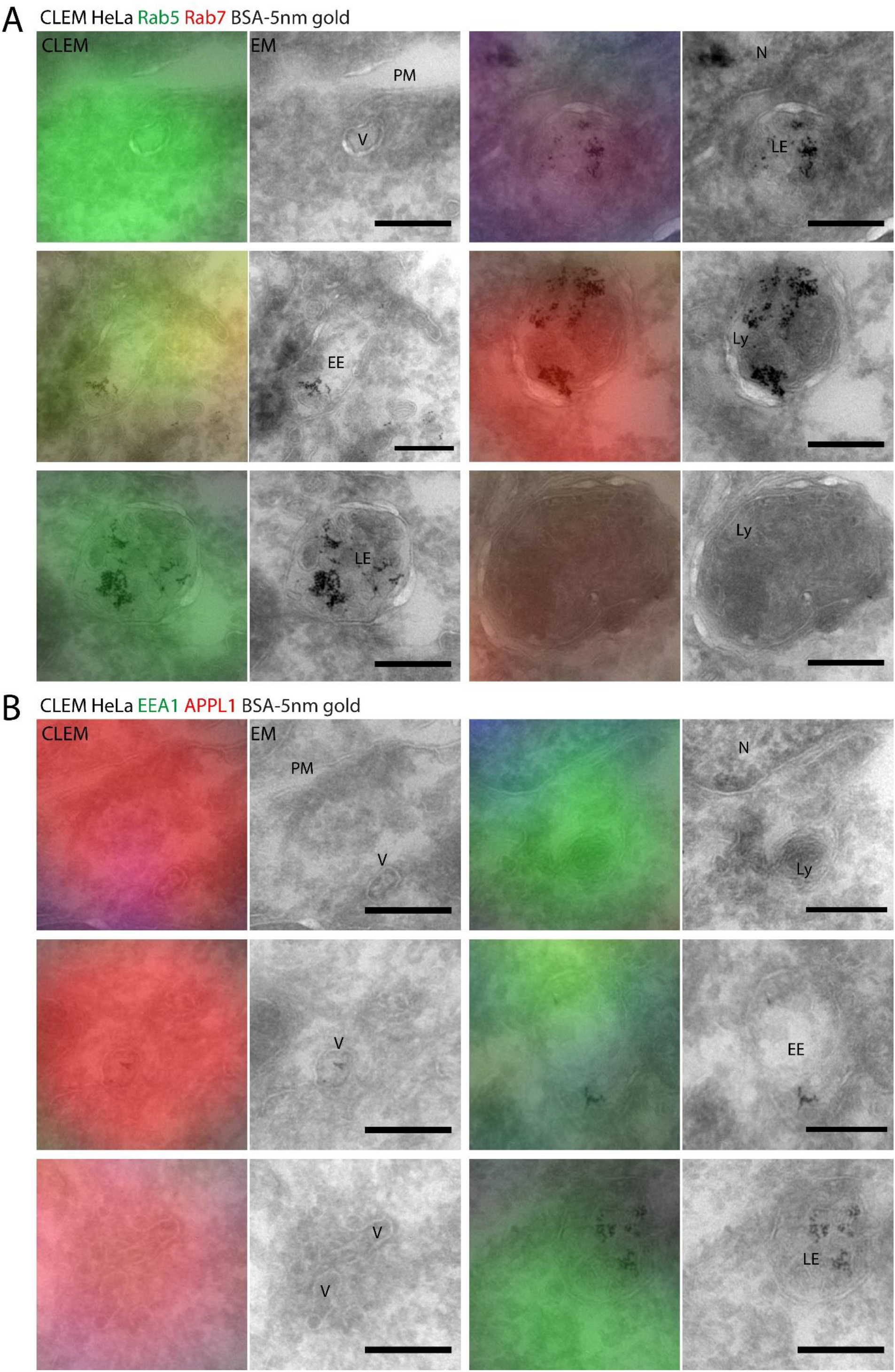
Original CLEM and EM images from pseudocolored examples. Preparation of samples as described in figures 3 and 4. A: original CLEM and EM images from figure 3C. B: CLEM and EM images of figure 4B. EE; early endosome, LE; late endosomes, Ly; lysosome, N; nucleus, PM; plasma membrane, V; vesicle. Scale bars 200nm.

**Figure S5.**
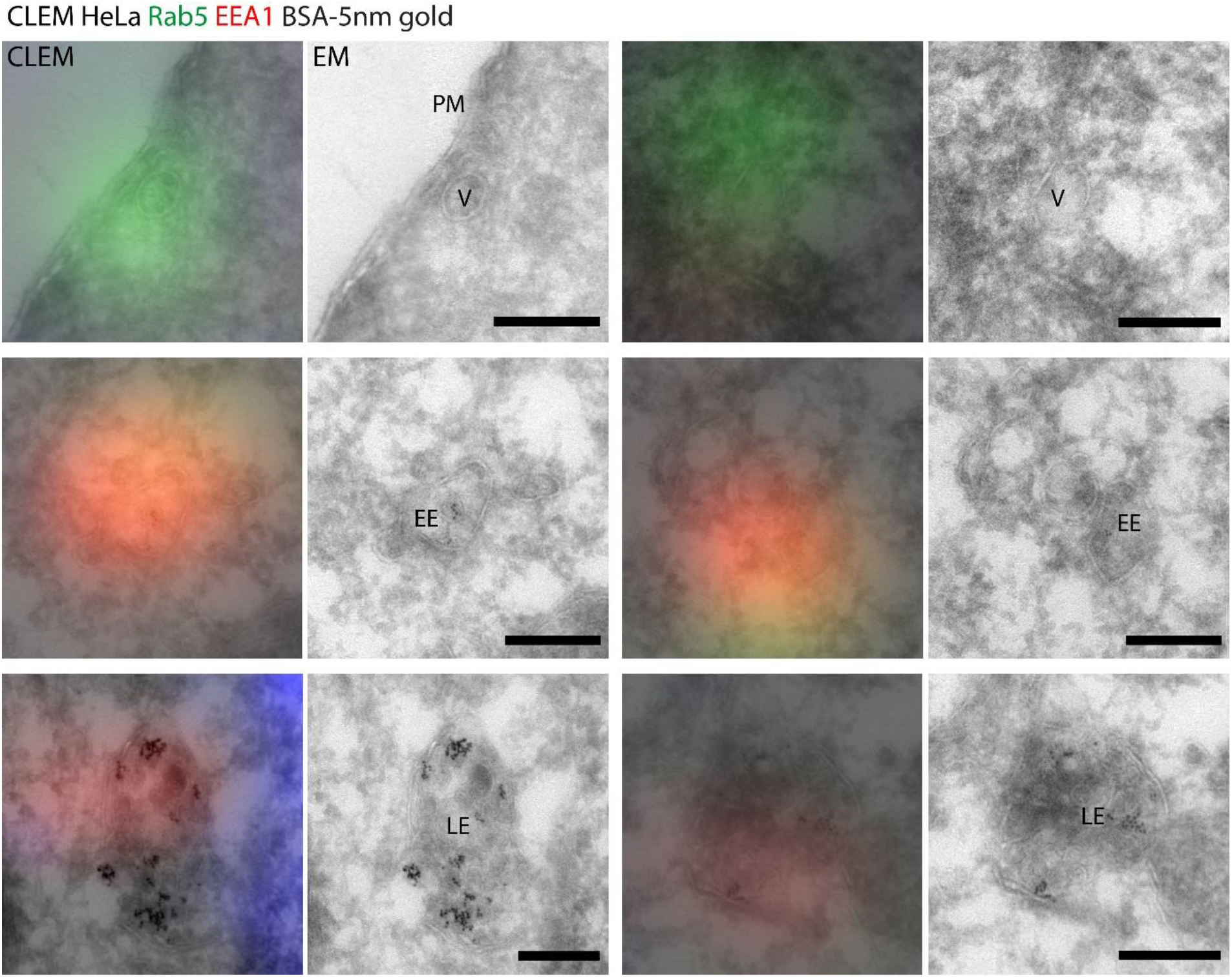
Original CLEM and EM images from pseudocolored examples in Figure 5A. Preparation of samples as described in figure 5. EE; early endosome, LE; late endosomes, PM; plasma membrane, V; vesicle. Scale bars 200nm.

**Figure S6.**
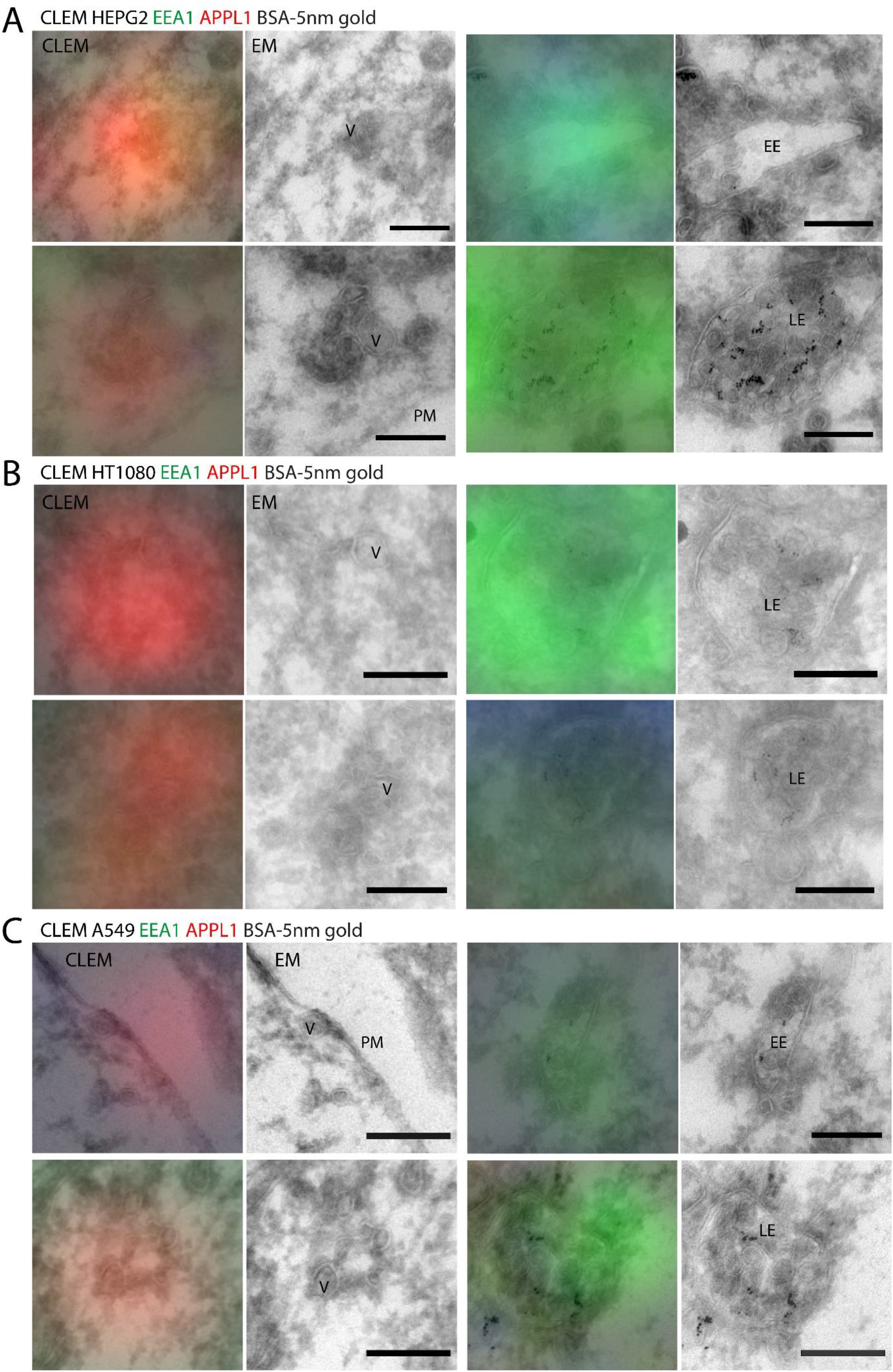
Original CLEM and EM images from pseudocolored examples. Preparation of samples as described in figure 6. A, B, C: CLEM and EM images of pseudocolored images in figure 6A, B and C, respectively. EE; early endosome, LE; late endosomes, PM; plasma membrane, V; vesicle. Scale bars 200nm.

**Table S1.** Organelle numbers, averages and standard deviation of quantification in F1. Cells from 2 independent replicates were analyzed. Values are Mean±SD.

| Combination (A:B) | Number of cells | Particles per cell for A | Particles per cell for B | Co-localized particles | % co-localized particles of A | % co-localized particles of B |
| --- | --- | --- | --- | --- | --- | --- |
| Rab5:Rab7 | 124 | 391.0±218.3 | 102.3±70.5 | 23.8±17.0 | 6.3±3.3 | 23.5±7.2 |
| CD63:Rab7 | 147 | 258.3±154.4 | 119.3±84.0 | 46.0±41.9 | 15.8±7.4 | 34.7±12.6 |
| CD63:CatD | 147 | 258.3±154.4 | 106.2±52.5 | 72.3±42.5 | 29.2±7.5 | 65.9±12.1 |
| Rab7:CatD | 147 | 119.3±84.0 | 106.2±52.5 | 27.3±21.9 | 22.0±7.8 | 23.9±12.0 |
| Rab5:APPL1 | 157 | 344.0±205.5 | 210.8±129.0 | 156.5±104.9 | 44.7±8.3 | 72.5±12.8 |
| Rab5:EEA1 | 133 | 342.4±164.5 | 121.2±59.9 | 90.4±49.0 | 26.4±7.5 | 73.5±13.6 |
| EEA1:APPL1 | 62 | 133.1±79.1 | 256.7±165.0 | 37.7±24.1 | 28.3±6.2 | 16.5±6.5 |

### Supplementary Note 1. CLEM method and notes

We here describe the on-section CLEM procedure and associated protocols in detail to aid other researchers in using this technique to its full advantage.

### Fixation

For most on-section CLEM approaches with immunolabelling, it is preferable to fix cells in 4% FA in 0.1 M phosphate buffer (PB) pH 7.4 without GA. Although addition of even small percentages of GA (0.05-0.2%) significantly improves the morphology, it has a negative effect on the efficacy of many antibodies. It additionally generates autofluorescent reaction products, which have to be quenched, e.g. with sodium borohydride (NaBH_4_), before labeling. Fixation with 4% FA is therefore preferable, the duration of the fixation can be optimized depending on the antibodies used. Generally speaking, longer fixation improves preservation of the ultrastructure while brief fixation is beneficial for labeling efficacy. We suggest testing durations in a range from 1 hour to overnight fixation. The performance of the antibody in regular Immuno-fluorescence (IF) can serve as an indicator for how well the antibody will work in CLEM and which fixation is best suited for it.

### Embedding, cryoprotection, and freezing

After fixation, cells are scraped, pelleted and embedded in 12% gelatin. Storage of the samples should be avoided for CLEM, as storage in low percentage FA (0.5-1.0%) will gradually progress the fixation, whilst storage in anything else will slowly undo the fixation and worsen preservation of the ultrastructure. Here follows a step-by-step protocol for sample preparation of adherent cells:

- Fixation using 4% FA in 0.1M PB pH 7.4 at selected duration
- Wash in PBS 3x
- Wash in PBS + 0.15% glycine for 10 minutes
- Replace with PBS
- Add pre-warmed (37°C) 2% gelatin 1:1 to PBS
- Scrape cells
- Transfer to microcentrifuge tube and pellet cells by centrifugation at 6000x *g* for 1 minute
- Remove supernatant
- Resuspend cells in pre-warmed (37°C) 12% gelatin
- Pellet cells by centrifugation at 6000x *g* for 1 minute
- Solidify gelatin on ice for 30 minutes
- Cut microcentrifuge tube above cell pellet
- Remove embedded cell pellet and cut into smaller blocks (∼1 mm^3^)
- Incubate in 2.3M sucrose overnight at 4°C, rotating end-over-end.
- Place blocks on pins suitable for use in cryo-microtomes
- Snap-freeze and store in liquid nitrogen

### Cryo-Sectioning

Embedded cells are sectioned in a cryo-ultramicrotome. The chamber, knife and specimen temperatures are set to -100°C. The blocks of gelatin-embedded cells are trimmed to a rectangular shape of ∼250 by 375 µm and cut to sections of 90-100 nm thickness. If ON 4% FA fixation is chosen, or GA is added, the temperature can be reduced to -110°C or -120°C to aid cutting thinner (70-90nm) sections. Section pickup is best done using a loop dipped in a 1:1 2.3M sucrose and 1.8% methylcellulose (MC) mixture. After pick-up, the sections are deposited on formvar and carbon-coated EM grids and can be stored 1-3 days at 4°C before use.

### Immuno-labelling and CLEM Workflow

#### Step-by-step protocol

- Place the grids sections down on PBS in a dish (e.g. a 3 cm Petri dish or multi-well plate) in a 37 °C stove for 20-30 minutes. This removes the sucrose + methylcellulose and the 12% gelatin in-between the cells
- Wash the grids on ∼100 µl drops of PBS + 0.15% Glycine at room temperature
- Wash in PBS 3 x 2 minutes
- Perform blocking step in PBS + 0.1% BSA-c + 0.5% cold fish-skin gelatin (FSG) for 10 minutes
- Incubate with primary antibody diluted in PBS + 0.1% BSA-c + 0.5% FSG for 60 minutes
- Wash in PBS + 0.1% BSA-c + 0.5% FSG 5 x 2 minutes
- Incubate with fluorescent secondary antibody and DAPI in PBS + 0.1% BSA-c + 0.5% FSG for 20+ minutes
- Wash in PBS + 0.1% BSA-c + 0.5% FSG 5 x 2 minutes
  - Optional: incubation with Protein A-Gold (PAG) for immuno-gold labeling, 20 minutes
  - Wash in PBS + 0.1% BSA-c + 0.5% FSG 5 x 2 minutes
- Wash in PBS 5 x 2 minutes
- Wash in demineralized H_2_O 3×2 minutes
- Submerge the grids in 50% glycerol in water
- Put grids on clean object slide, sections facing up
- Add a small droplet of glycerol
- Cover with a clean coverslip
- Perform imaging of the sections on a suitable fluorescence microscope
- Remove the oil carefully from the coverslip, e.g. with EtOH
- Unmount the grids in distilled water
- Carefully dry the backside of the grids
- Place grids on drops of H_2_0
- Fix in PBS + 1% GA for 5 minutes
- Wash in H_2_O 8 x 2 minutes
- Postfix in 2% UA pH7 for 5 minutes
- Wash 2x in MC/UA pH4 on ice briefly
- Incubate with MC/UA pH4 on ice for 5-10 minutes
- Loop out with a metal ring and filter paper. This forms a thin layer of MC on the sections.
- Dry the grid

#### Notes

The IF labeling can slowly deteriorate while the grids are kept in 50% glycerol. It is therefore advised to only process and image a few grids at a time, keeping the time between secondary labeling and fixation in 1% GA to 30-45 minutes. The remaining grids can be left in the secondary labeling step or on Hoechst solution until they are ready for processing and imaging. Take care to fully submerge the grids in 50% glycerol, since air trapped between the grids and the coverslip can break the formvar film. The slide- and coverglass used to sandwich the grid should be thoroughly cleaned beforehand. Use the ‘squeaky clean coverslips’ protocol described by Waterman-Storer, C. M. ^103^. When retrieving the grids from between the slide- and coverglass, be careful not to mix the immersion oil from the microscope with the 50% glycerol, as this might leave oil on the sections. Drying of the backside of the grids should be performed with care, as complete drying of the grid will result in deterioration of the ultrastructure. For fluorescence imaging, we recommend a widefield microscope with a fast, automated stage, a sensitive camera and the ability to make image tilesets. Use a 60-100x oil objective to create high-resolution image tilesets of (parts of) the ribbon of sections. Having a large area imaged in the fluorescence microscope will aid finding and selecting areas to image with EM. On slower microscopes or larger samples, it is also possible to create a tileset at a lower magnification as an overview, and then select areas of interest for imaging using a higher magnification objective.

### Correlation

To select the area to be imaged in EM, begin by finding a region that has been imaged in fluorescence microscopy. Select an area that has well-preserved ultrastructure in EM and strong, in-focus signal in fluorescence microscopy. For EM imaging, we recommend using a TEM with high image quality at 30,000-100,000x magnification capable of generating image tilesets. Of the selected area, generate a tileset at 40,000-50,000x magnification of roughly 500-1000 µm^2^. After postprocessing of the data to generate a large, continuous EM image, overlay the EM and fluorescence images using software designed for this operation such as the ec-CLEM plug-in in Icy, or by manually translating the images in suitable software (such as Adobe Photoshop). In either case, use reference points that are not the primary object of study and are available in both EM and fluorescence to overlay your images. These can be edges of nuclei, cell outlines, mitochondria or specific bimodal probes (fiducial markers), as long as an appropriate marker has been included in the labeling procedure.

### Reagents

- **Phosphate Buffer** (**PB**) 0.2M: Make by mixing 0.2M NaH_2_PO_4_.1H_2_O and 0.2M Na_2_PO_4_.2H_2_O in 19:81 ratio. Dilute as needed.
- **2.3M sucrose:** Sucrose D(+) saccharose in 0.1M PB.
- **UA PH7**: Uranyloxalicacetate pH7. Dissolve 4g uranylacetate in 100ml H_2_O. Dissolve 3.8g oxalic acid in 100ml H_2_O. Mix 1:1 and add NH_4_OH until pH7.0 is reached. Use pH indicator sticks. Filter 0.45 µm before use.
- **MC/UA pH4**: Dissolve 0.4g Uranylacetate in 10ml H_2_O. Mix 1:9 with 2% cellulose in H_2_O.
- **MC**: 1.8% Methylcellulose (25 centipoises, Sigma M-6385) in H_2_O.
- **Gelatin**: For 12%, add 12g food-grade gelatin to 75ml 0.1M PB. Warm to 60°C and stir. Add 100µl 20% Na-Azide, add 0.1M PB up to 100ml. Dilute in 0.1M PB as needed.
- **FA formaldehyde**: For 16% stock aliquots. Add 80g PFA (prilled, Sigma-Aldrich 441244) to 400ml H_2_O. Warm to 60°C and stir for 15min. Add 0.1M NaOH until pH is 7, use indicator sticks. Stir for 30min at 60°C. Cool to RT, check pH again. Add H_2_O up to 500ml. Filter solution. Freeze aliquots. Thaw for use, sometimes heating is required. Do not use if solution does not turn clear.
- **BSA**: Bovine serum albumin fraction V (Sigma A-9647). Dilute in H_2_O.
- **PAG**: Protein-A gold. Protein-A conjugated to colloidal gold particles. Made in-house, Cell Microscopy Core, UMC Utrecht. Available online.

